# Unraveling the Mechanism of HIV-1 Hypersusceptibility to Tenofovir Imparted by Islatravir Resistance Mutations

**DOI:** 10.64898/2026.06.02.725532

**Authors:** Shreya M. Ravichandran, Alexa A. Snyder, Katelyn H. Kim, Isabella L. Kaufman, Xin Wen, Eleftherios Michailidis, Karen A. Kirby, Stefan G. Sarafianos

## Abstract

In response to the newly approved antiretroviral therapy (ART) islatravir (ISL), the M184V and A114S resistance mutations have emerged in the human immunodeficiency virus type 1 reverse transcriptase (HIV-1 RT). These mutations markedly hypersensitize RT to the globally administered ART tenofovir disoproxil fumarate (TDF). We have solved six structures – four by X-ray crystallography and two by cryo-EM – that capture the single- and double-mutant RTs during inhibitor incorporation and demonstrate the role of the mutations in altering protein-antiviral interactions. These snapshots reveal that the smaller, more flexible TDF diphosphate (TDF-DP) can better accommodate mutation-induced active site changes than ISL triphosphate (ISL-TP). Structural differences between the two inhibitors are consistent with biochemical determination of inhibitory constants (*K*_i_s), highlighting differences at the step of inhibitor incorporation. Virological evaluation of ISL and TDF combinations reveals additive inhibition of HIV-1. Given the converse ISL hypersusceptibility imparted by the TDF-resistant K65R mutation, we propose ISL and TDF as a combination that can inspire future therapeutic options.

## INTRODUCTION

With over 40 million cases of human immunodeficiency virus (HIV) globally as of 2024, developments toward novel antiretroviral therapies (ARTs) persist worldwide. HIV type 1 (HIV-1) is largely incurable^1–3^, leading to lifelong-administered ARTs typically recommended as daily regimens, with long-acting options growing in prevalence^4, 5^. Consequently, ART-resistant viral mutations can emerge decrease the effectiveness of ARTs^6, 7^.

ARTs are typically administered in combinations of two-to-three antivirals that synchronously inhibit multiple steps of the HIV replication cycle, the most common target being the HIV-1 reverse transcriptase (RT)^8^. HIV-1 RT is crucially responsible for synthesizing proviral, double-stranded DNA (dsDNA), templated by the negative-sense, single-stranded RNA (ssRNA) genome packaged within an infectious particle^9^. This provirus can then be integrated into the infected host cell’s genome and the immune cell’s machinery hijacked for future rounds of viral replication. The key activities of HIV-1 RT have resultingly prompted the U.S. Food and Drug Administration (FDA)’s approval of three ART classes against the enzyme: nucleoside reverse transcriptase inhibitors (NRTIs), nucleoside reverse transcriptase translocation inhibitors (NRTTIs), and non-nucleoside reverse transcriptase inhibitors (NNRTIs)^8, 10–14^. Notably, all first-line ART combinations to-date have included at least one RT-targeting drug^8^, and the effectiveness of targeting RT is used in long-acting ARTs and PrEP regimens, too^4, 5, 15^.

Tenofovir (TDF; Gilead Sciences, Inc.) is a deoxyadenosine analog that is routinely administered to a large proportion of people living with HIV worldwide^16, 17^, its utility expanding to people living with Hepatitis B virus (HBV) as well^18, 19^. As an NRTI, TDF lacks a 3’-OH group (Fig. S1A), the basis of its sole inhibition modality: obligate chain termination (OCT)^20^. Here, NRTI incorporation into the nascent proviral DNA prevents a subsequent nucleotide from successfully forming a phosphodiester bond with the 3’-OH-lacking analog. Consequently, OCT stops provirus synthesis, preventing HIV-1 from elongating the sequence that would otherwise translate into future infectious particles. Also of note, as an acyclic inhibitor, TDF is unique among NRTIs for its pointed lack of a deoxyribose ring all together (Fig. S1A), giving rise to an inherent flexibility.

Islatravir (ISL; Merck & Co., Inc.) is the first-in-class NRTTI^10, 11^, a deoxyadenosine analog approved in April 2026 for use alongside doravirine (DOR) as the newest, once-daily ART, IDVYNSO™ (Merck & Co., Inc.)^13, 14, 21^. Interestingly, NRTTIs retain the 3’-OH group found in naturally occurring nucleotides, though they also possess a unique 4’-ethynyl (4’-E) group (Fig. S1A), a hydrophobic moiety that bolsters the ligand’s interaction with a conserved hydrophobic pocket at the polymerase active site^12^. Effectively, NRTTIs inhibit differentially from NRTIs by acting either through OCT or delayed chain termination (DCT) through its 3’-OH group ^11^.

Previously, clinical findings have reported that ISL treatment leads to the emergence of ART-resistant HIV-1 RT mutants, including RT_M184V_^22, 23^. Historically, M184V has been associated with high-level emtricitabine (FTC)/lamivudine (3TC) resistance and azidothymidine (AZT) hypersusceptibility^24–27^. The modest ISL resistance conferred by M184V is heightened, though, by simultaneous presence of the ISL-unique A114S mutation at the polymerase active site, as we and others reported^22, 23^. Despite ISL’s remarkably high potency, establishment of a new targeting mechanism, and robust long-acting potential^28^, the selection for HIV-1 RT_M184V_, RT_M184V/A114S_, and additional mutants underscores the need to characterize and overcome mechanism(s) of resistance. Interestingly, previous *in cellulo* works have suggested that RT_M184V/A114S_ hypersensitizes – or increases susceptibility of – HIV-1 to the TDF prodrug^29^. This is unexpected given that both ISL and TDF are deoxyadenosine analogs, and the mechanisms by which the M184V and A114S mutations cause ISL resistance and TDF hypersusceptibility remain largely unknown, though recent studies have emerged^30^. As a result, our work utilizes biochemical, virological, and structural studies of these mutant-antiviral combinations to explain these contrasting profiles and inspire novel treatment approaches.

In the current HIV-1 therapeutics era, drug resistance is a driver of novel treatment strategies. Accordingly, ISL’s recent clinical introduction and TDF’s longstanding status as one of the most-prescribed ARTs worldwide provide the foundations for an unexplored, promising pair addressing drug resistance concerns. Furthermore, given their TDF hypersusceptibility, emergence of RT_M184V_ and RT_M184V/A114S_ motivates the addition of TDF to ISL treatment for people living with or at risk of experiencing these ISL-resistant mutants. From the summation of our findings, we therefore highlight ISL and TDF as a potential combination of two dATP analogs that can combat increasingly prevalent, drug-resistant HIV-1 infections worldwide.

## RESULTS

### Structures capturing incoming ISL-TP reveal a “steric crowding” effect at the active site

We began by determining six molecular structures of complexes trapped in the process of incorporating inhibitor into the DNA chain (Tables S1, S2). Each ternary complex comprised HIV-1 RT, dsDNA, and antiviral, varying by the RT mutant-in-question (RT_M184V/A114S_, RT_M184V_, RT_A114S_) and the phosphorylated antiviral-of-interest (ISL-TP or TDF-DP). We crosslinked HIV-1 RT residue Q258C to a thioalkyl group on a modified guanosine within the P_d18_* sequence and terminated the primer with ddGMP to prevent the incoming ISL-TP or TDF-DP from covalently incorporating into the nascent strand. In this frozen state, we could identify the mutation- and antiviral-dependent changes implicated in ISL-TP resistance and/or TDF-DP hypersusceptibility during inhibitor incorporation.

We were able to successfully capture ISL-TP or TDF-DP residing at the nucleotide binding (*N*) site in all six structures (Fig. S4). Across structures, we identified the “sandwiching” of the inhibitor between the 114 and 184 residues. This apparent sandwiching of the 4’-E involves interactions between the inflexible, β-branched V184 – positioned within 3.3 Å from the 4’-E of ISL-TP – and, from the other side, an S114-stabilized network of interactions involving several metal-coordinated residues. S114 uniquely protrudes into the primary Mg^2+^ coordination site required for successful polymerization of the nascent DNA chain (Fig. 1A). Focusing on RT_M184V/A114S_ (PDB ID 12SK; 2.79 Å), we observed no significant changes to the V184 backbone (Fig. 1B). The increased steric crowding appeared compounded by the S114 side chain seemingly pushing both the C_α_ of G112 and the Mg^2+^-coordinated carbonyl of V111 (Fig. 1C). The carbonyl of V111 itself did not show a major difference in distance to the metal. Additionally, S114 seemingly caused the D110 side chain to rotate orthogonally (Fig. 1C). In the new rotamer state, D110 was 2.7 Å from the metal as opposed to its original 2.0-Å distance in RT_WT_ (PDB ID 5J2M^12^). Further observations of the catalytic triad (D110, D185, D186) revealed that S114 sat nearly one Å closer to D185 than A114 and at an angle conducive to an H-bond interaction, locking the 4’-E between this region and V184 (Fig. 1D). Therefore, we hypothesized that RT_M184V/A114S_ confers resistance to ISL-TP through strong steric crowding caused by both V184 and S114.

**Fig. 1.**
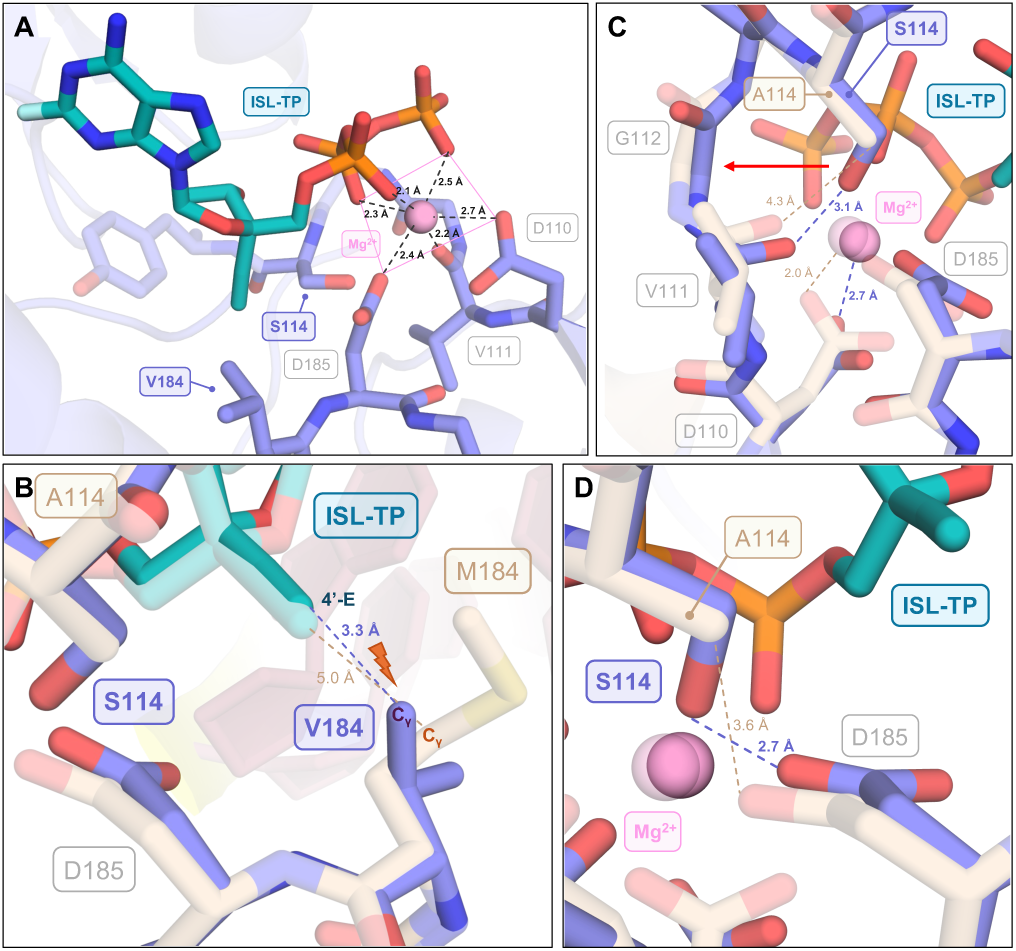
Crystal structure of HIV-1 RT_M184V/A114S_ with dsDNA_ddGMP_ and incoming ISL-TP. *[A]* The polymerase active site magnesium geometry is displayed, with the M184V and A114S mutations labeled. *[B]* Overlay of RT_WT_ (PDB ID 5J2M) and RT_M184V/A114S_ (PDB ID 12SK) at residue 184. *[C]* Comparison of RT_WT_ (PDB ID 5J2M) and RT_M184V/A114S_ (PDB ID 12SK) around residues 110-114. *[D]* Overlay of RT_WT_ (PDB ID 5J2M) and RT_M184V/A114S_ (PDB ID 12SK) near residue 185. PyMOL color palette: 12SK (RT_M184V/A114S_): purple; 5J2M (RT_WT_): tan; dsDNA backbone: yellow; dsDNA bases: pink; ISL-TP: teal; Mg^2+^ ion: light pink; phosphorus: orange; oxygen: red; nitrogen: blue; sulfur: yellow; WT *vs.* mutant residue labels: wheat *vs.* slate; addl. residue labels: gray. Figure created in PyMOL v3.1.3 (Schrödinger, LLC.).

### Structures of incoming TDF-DP reveal beneficial mutation-driven interactions at the active site

Our remaining ternary complexes utilized TDF-DP as the incoming, non-covalently bound nucleotide. In assessing the double-mutant structure (PDB ID 12SW; 3.20 Å), we first demonstrated that there were no obvious perturbations to the expected metal ion coordination at the polymerase active site (Fig. 2A). We then compared mutation-driven interactions of TDF-DP within RT_M184V/A114S_ to those of TDF-DP associated with RT_WT_ (PDB ID 1T05^31^). Visibly, there appeared to be a sterics-related benefit to substituting V184 in-place of the methionine, for the distance of the nearest V184 C_γ_ to the analog’s C_2’_ was further away than the original ∼5-Å distance of the M184 C_γ_ (Fig. 2B). Additionally, there was an extension of the S114 side chain in the direction of the primary Mg^2+^ coordination scheme, but we did not identify concomitant changes to the D110 rotamer as we did with ISL-TP (Fig. 2A). Instead, we noticed that S114 was positioned 2.6 Å from the TDF-DP β-phosphate at a distance and angle consistent with an H-bond interaction (Fig. 2C). Taken together, these data are consistent with the reported hypersusceptibility of the double-mutant RT to TDF-DP.

**Fig. 2.**
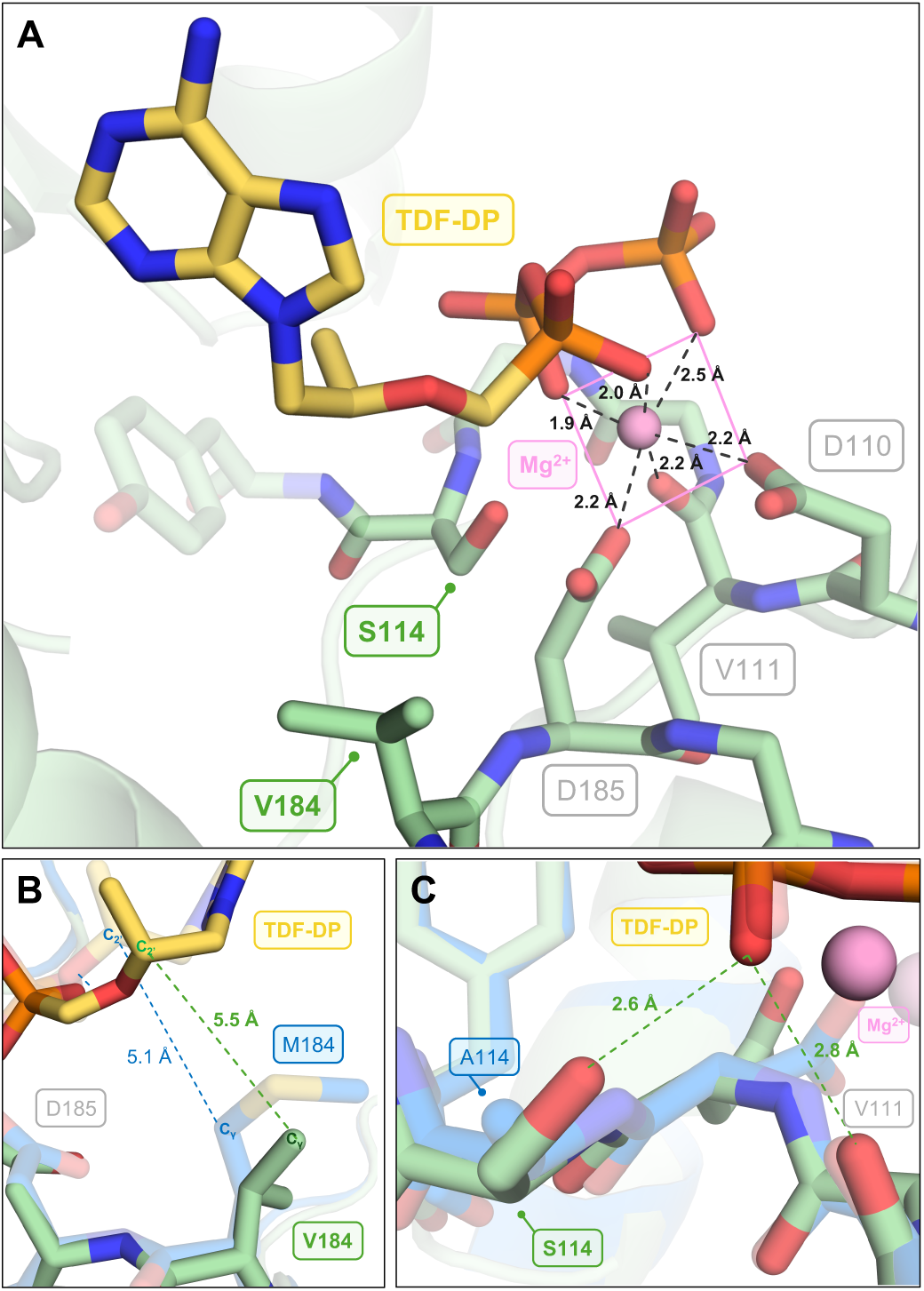
Crystal structure of HIV-1 RT_M184V/A114S_ with dsDNA_ddGMP_ and incoming TDF-DP. *[A]* The polymerase active site magnesium geometry is displayed, with the M184V and A114S mutations labeled. *[B]* Overlay of RT_WT_ (PDB ID 1T05) and RT_M184V/A114S_ (PDB ID 12SW) at residue 184. *[C]* Comparison of RT_WT_ (PDB ID 1T05) and RT_M184V/A114S_ (PDB ID 12SW) near the TDF-DP β-phosphate. PyMOL color palette: 12SW (RT): light green; 1T05 (RT): blue; TDF-DP: yellow; Mg^2+^ ion: pink; phosphorus: orange; oxygen: red; nitrogen: blue; sulfur: yellow; WT *vs.* mutant residue labels: blue *vs.* light green; addl. residue labels: gray. Figure created in PyMOL v3.1.3 (Schrödinger, LLC.).

### RT_M184V/A114S_ is highly ISL-TP incorporation-deficient due to a severe loss of substrate affinity

We assessed if inhibition by ISL-TP or TDF-DP of *in vitro* primer extension along a 31-mer template was affected by the mutation(s) present (Table S3). Our findings revealed that RT_M184V/A114S_ exhibited ∼116-fold ISL-TP resistance and 25.8-fold TDF-DP hypersusceptibility in gel-based DNA extension assays (Fig. 3). Both RT_M184V_ and RT_A114S_ showed intermediate ISL-TP resistance and TDF-DP hypersusceptibility (Fig. 3). Therefore, these data demonstrate that HIV-1 RT_M184V/A114S_ is an ISL-TP-resistant and TDF-DP-hypersusceptible enzyme.

**Fig. 3.**
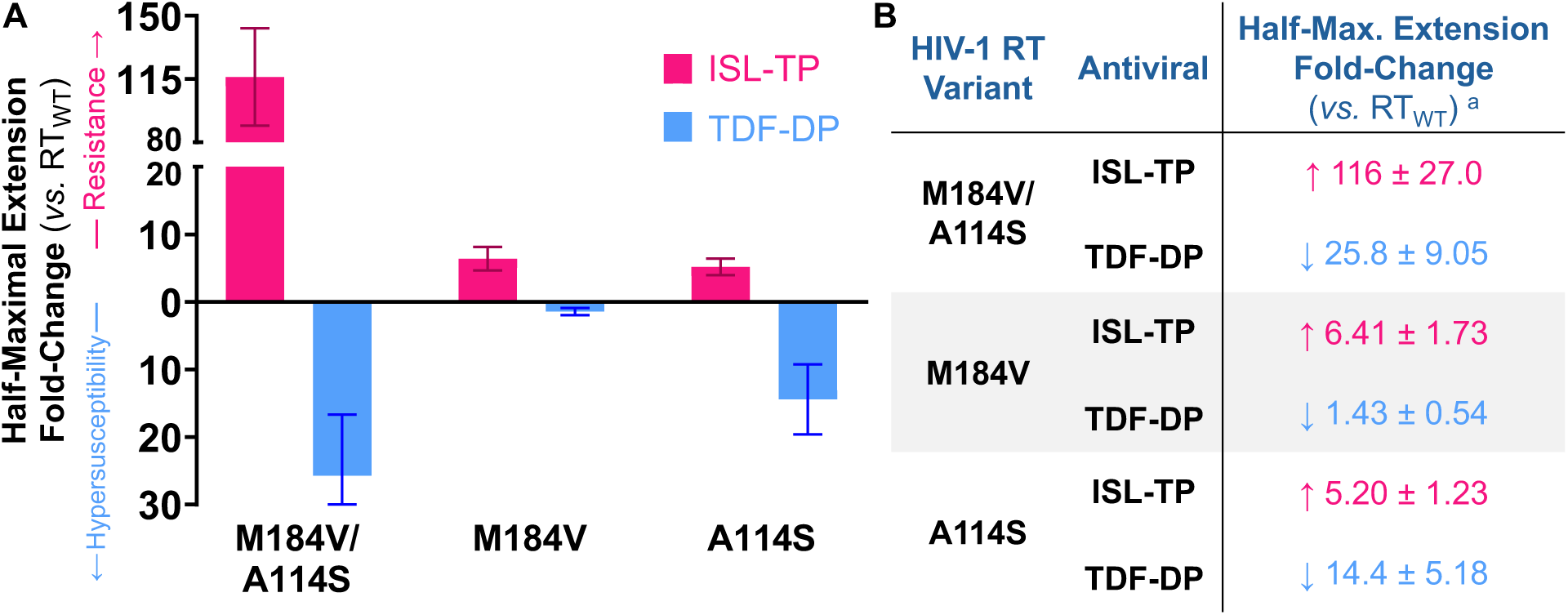
ISL-TP and TDF-DP profiles among HIV-1 RTs during *in vitro* primer extension. *[A-B]* Graph created using GraphPad Prism 9. Fold-changes in antiviral concentrations leading to half-maximal DNA extension are reported among the antiviral-RT combinations. Values reported from three experiments. *[a]* Fold-change values are provided with propagated standard errors as described in Materials and Methods. Color palette: resistance: pink, upward-facing arrow; hypersusceptibility: blue, downward-facing arrow. Graph and non-linear regression calculations using four-parameter inhibition done in GraphPad Prism 9.

We then assessed each step of reverse transcription with RT_WT_, RT_M184V/A114S_, RT_M184V_, and RT_A114S_ (Fig. S1B). We initially conducted time-courses of dATP, ISL-TP, and TDF-DP incorporation to locate a timeframe with a constant nucleotide incorporation rate. Though we did not seek immediate conclusions from this experiment, there was an evident difficulty in the ability of RT_M184V/A114S_ to incorporate ISL-TP over time (Fig. S6). Moreover, we observed a linear incorporation rate across the 12 mutant/nucleotide combinations intersecting at the three-min timepoint, therefore selecting this reaction length to test steady-state inhibitor incorporation.

We used nucleotide concentration-dependent reactions to reveal the steady-state kinetics (*K*_m_, *k*_cat_, *k*_cat_/*K*_m_) associated with dATP, ISL-TP, and TDF-DP incorporation. Firstly, we checked the relative fitness of each RT through the incorporation kinetics of the natural substrate, dATP. Compared to RT_WT_, we saw a moderate decrease in dATP incorporation efficiencies in RT_M184V/A114S_ and RT_A114S_ (Table 1, Fig. 4). In then gauging nucleotide incorporation by RT_WT_, we identified the highest selectivity index (SI) associated with ISL-TP (1.26X) yet a relatively more-unfavorable SI for TDF-DP (0.20X) (Table 1). Strikingly, RT_M184V/A114S_ displayed less-than-one-hundredth of RT_WT_’s ISL-TP incorporation efficiency (Table 1, Fig. 4). We determined this to arise from a 240-fold growth in the associated *K*_m_ – from 180 nM (RT_WT_) to 43 µM – and a 20% decrease in the *k*_cat_, dropping the resulting catalytic efficiency to 0.04 min^-1^ µM^-1^ (Table 1, Fig. 4). In comparison, each of the single mutants showed slightly deficient ISL-TP incorporation compared to RT_WT_, though they were substantially more efficient than RT_M184V/A114S_ (Table 1).

**Fig. 4.**
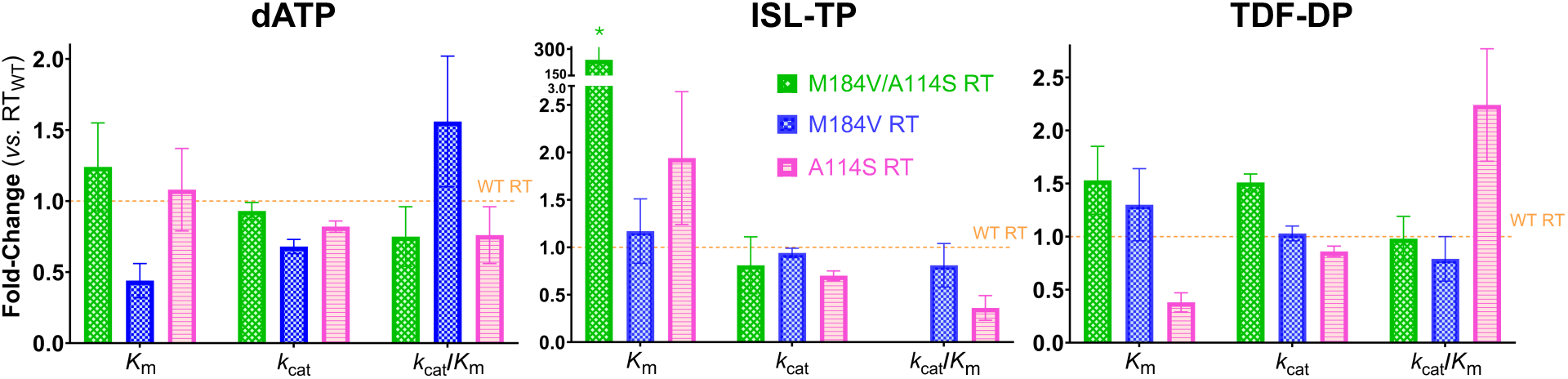
Fold-changes in inhibitor incorporation kinetics among HIV-1 RT mutants. Comparisons of *K*_m_, *k*_cat,_ and *k*_cat_/*K*_m_ are reported between HIV-1 RT variants among nucleotides. All fold-changes are graphed with propagated standard error bars as described in Materials and Methods. Values calculated from results of three-to-five experiments. Color palette: RT_WT_: orange line; RT_M184V/A114S_: green bar; RT_M184V_: blue bar; RT_A114S_: pink bar. Asterisk (*) denotes y-axis break between 3.0X – 150X. Graphs created in GraphPad Prism 9.

**Table 1.** Impact of the M184V and A114S HIV-1 RT mutations on ISL-TP and TDF-DP incorporation. Under steady-state conditions, fold-changes in the *K*_m_, *k*_cat_, and catalytic efficiency alongside selectivity indices are reported for nucleotide-RT pairings. Using the Cheng-Prusoff equation, fold-changes in *K*_i_s were determined. Values calculated from results of three-to-five experiments. *[A]* All values are reported with propagated standard errors calculated as described in Materials and Methods. Non-linear regression calculations performed under “Michaelis-Menten Kinetics” in GraphPad Prism 9. *[B]* Cheng-Prusoff equation: *K*_i_ = IC_50_ / (1 + [S]/*K*_m_); S = dATP; “IC_50_s” and [S] were taken from *in vitro* primer extension assays; dATP *K*_m_s were taken from single-nucleotide incorporation assays. Color palette: resistance: pink, upward-facing arrow; hypersusceptibility: blue, downward-facing arrow.

| HIV-1 RT Variant | dNTP | $K_m$ Fold-Change<br>(vs. $RT_{WT}$)<sup>A</sup> | $k_{cat}$ Fold-Change<br>(vs. $RT_{WT}$) | $k_{cat}/K_m$ Fold-Change<br>(vs. $RT_{WT}$) | Selectivity Index<br>(vs. dATP) | $K_i$<br>($\mu M$) | $K_i$ Fold-Change<br>(vs. $RT_{WT}$)<sup>B</sup> |
| --- | --- | --- | --- | --- | --- | --- | --- |
| Wild-Type | dATP | - | - | - | - | - | - |
| | ISL-TP | - | - | - | 1.26 ± 0.36 | 0.05 ± 0.02 | - |
| | TDF-DP | - | - | - | 0.20 ± 0.05 | 6.59 ± 3.41 | - |
| M184V/A114S | dATP | 1.24 ± 0.31 | 0.93 ± 0.06 | 0.75 ± 0.21 | - | - | - |
| | ISL-TP | 239 ± 144 | 0.81 ± 0.30 | <0.01 ± <0.01 | <0.01 ± 0.02 | 6.49 ± 1.62 | ↑ 143 ± 68.9 |
| | TDF-DP | 1.53 ± 0.32 | 1.51 ± 0.08 | 0.98 ± 0.21 | 0.26 ± 0.07 | 0.32 ± 0.13 | ↓ 20.9 ± 14.0 |
| M184V | dATP | 0.44 ± 0.12 | 0.68 ± 0.05 | 1.56 ± 0.46 | - | - | - |
| | ISL-TP | 1.17 ± 0.34 | 0.94 ± 0.05 | 0.81 ± 0.23 | 0.65 ± 0.19 | 0.13 ± 0.04 | ↑ 2.85 ± 1.52 |
| | TDF-DP | 1.30 ± 0.34 | 1.03 ± 0.07 | 0.79 ± 0.21 | 0.10 ± 0.03 | 2.05 ± 0.99 | ↓ 3.22 ± 2.28 |
| A114S | dATP | 1.08 ± 0.29 | 0.82 ± 0.04 | 0.76 ± 0.20 | - | - | - |
| | ISL-TP | 1.94 ± 0.70 | 0.70 ± 0.05 | 0.36 ± 0.13 | 0.60 ± 0.20 | 0.25 ± 0.07 | ↑ 5.59 ± 2.72 |
| | TDF-DP | 0.38 ± 0.09 | 0.86 ± 0.05 | 2.24 ± 0.53 | 0.59 ± 0.15 | 0.49 ± 0.22 | ↓ 13.3 ± 9.03 |

In contrast to ISL-TP, RT_M184V/A114S_ carried out relatively unaffected TDF-DP incorporation, evidenced by no change in enzymatic efficiency (Table 1). We identified the unchanged efficiency as the effect of similar fold-increases in the associated *K*_m_ (1.5X) and *k*_cat_ (1.5X) *vs.* those of RT_WT_ (Table 1, Fig. 4). We additionally observed an improved TDF-DP incorporation efficiency with RT_A114S_ alone (2.2X) that primarily stemmed from a decrease in the *K*_m_ (Table 1, Fig. 4). Overall, our tests of single-nucleotide incorporation across all RT-antiviral combinations revealed that RT_M184V/A114S_ exhibits ISL-TP resistance due to a significant loss of substrate affinity during nucleotide incorporation into the nascent chain, with contributions from both incorporation-deficient mutations. At the same time, we find that RT_M184V/A114S_ does not cause strong changes to the kinetics governing TDF-DP incorporation.

### Inhibitory constants depict changes to the competitive abilities of the inhibitors *vs.* dATP

Due to the apparently unaffected efficiency of RT_M184V/A114S_ in incorporating TDF-DP, we assessed the inhibitory constant (*K*_i_) of each inhibitor against the mutants, now with the presence of dATP taken into consideration. In doing so, we evaluated whether the amount of ISL-TP or TDF-DP needed to *[1]* overcome the presence of the natural substrate and then *[2]* bind the target site would change depending on the RT being targeted. To calculate the *K*_i_ of a competitive inhibitor under non-cooperative assumptions^32^, we applied the Cheng-Prusoff equation, used the steady-state dATP incorporation-derived *K*_m_ values (Table S4), and consulted the primer extension-derived half-maximal inhibition values as stand-in “IC_50_s” obtained in the presence of 10 μM dATP (Fig. S5). Resultingly, we observed a 21-fold improvement in the *K*_i_ of TDF-DP with RT_M184V/A114S_ *vs.* RT_WT_ (Table 1). Opposingly, the *K*_i_ of ISL-TP with the double-mutant worsened by 143-fold *vs.* WT, consistent with the previously identified impairment to ISL-TP incorporation kinetics (Table 1). Thus, the improved *K*_i_ for the double-mutant over RT_WT_ in the presence of dATP leads to the reported RT_M184V/A114S_ hypersusceptibility toward TDF-DP.

### RT_M184V/A114S_ translocation is largely unchanged on inhibitor-terminated nucleic acid substrate

Following steady-state incorporation, we evaluated whether the translocation ability of each RT along inhibitor-terminated DNA would differ based on inhibitor (Table S3). After terminating DNA primer with each dATP analog (ddATP, ISL-TP, TDF-DP), we performed hydroxyl radical DNA footprinting in the absence of incoming dTTP (Fig. S7A)^33^. In comparing post-translocated, primer-binding (*P) vs.* pre-translocated, nucleotide binding (*N)* RT complexes (Fig. S10), roughly 60% of all RTs translocated with ddAMP as the primer-terminating base (Fig. S7A, Table S5). In contrast, ISL-MP and TDF complexes remained primarily pre-translocated (Fig. S7A). For both antivirals, RT_M184V_ marginally showed the most movement, with 29.8% of ISL-MP and 26.3% of TDF *P* complexes (Table S5). Nonetheless, Benjamini-Hochberg (BH)-corrected t-tests at *Q* = 0.01 in each nucleotide condition showed insignificant differences in *P* complex levels without added dTTP (Fig. S7A).

At concentrations between 0-750 µM dTTP, ddAMP-associated RTs facilely translocated up to 90% (Fig. 5A). Over half of all ISL-MP complexes translocated up to 80% at high dTTP concentrations, yet RT_M184V_ required minimal dTTP unlike the other RTs (Fig. 5A). With TDF, we found that RT_M184V/A114S_ and RT_A114S_ were less able to translocate in the presence of lower dTTP concentrations, only half as much as RT_WT_ or RT_M184V_ up to 200 µM dTTP (Fig. 5A). We propose that the subtle difference in the double-mutant’s translocation efficiency along TDF-terminated primer may provide a minor TDF hypersusceptibility mechanism by forming a small population of immobile RTs, though differences may not be meaningful at cellular dTTP levels. Thus, we find that RT translocation does not significantly contribute to ISL resistance or TDF hypersusceptibility.

**Fig. 5.**
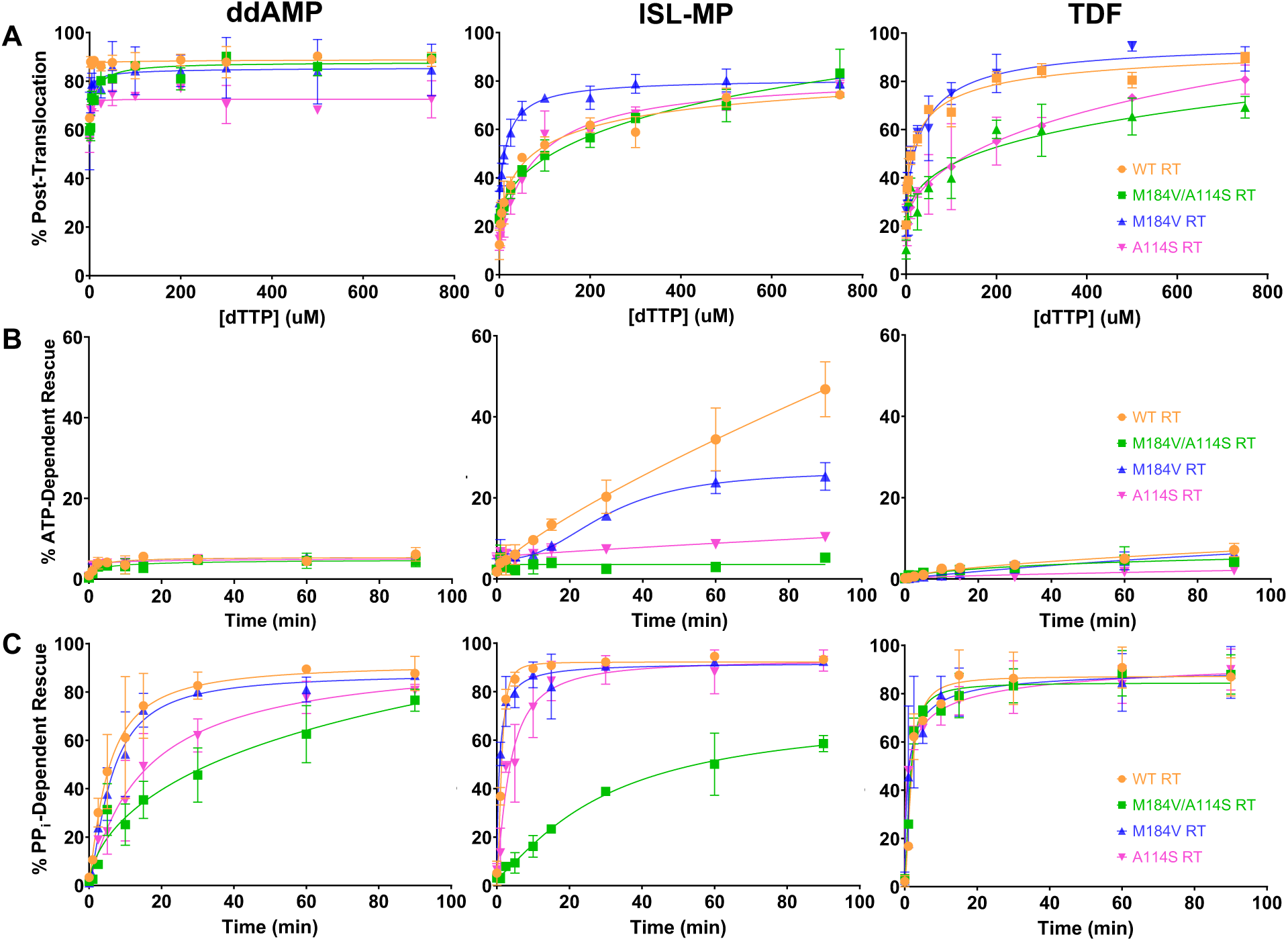
Impact of M184V and A114S mutations on HIV-1 RT translocation and inhibitor excision / DNA rescue. *[A]* Along a concentration gradient of incoming dTTP, the prevalence of translocation is reported among the RTs based on the chain-terminating analog. *[B]* ATP-mediated excision was tested along a 90-min time course in the presence of 3.5 mM ATP. *[C]* PP_i_ -mediated excision was tested along a 90-min time course in the presence of 150 µM PP_i_. Values reported from two-to-six experiments. Color palette: RT_WT_: orange; RT_M184V/A114S_: green; RT_M184V_: blue; RT_A114S_: pink. Graphs created in GraphPad Prism 9.

### ATP and PP_i_-mediated primer unblocking are overall unchanged among RT variants

We assessed the ability of each RT to carry out a two-step resistance mechanism: *[1]* “unblocking”: ATP- and pyrophosphate (PP_i_)-mediated inhibitor excision, followed by *[2]* “rescue”: limited DNA extension using dATP, dTTP, and ddGTP^34^. We purified DNA terminated with ddAMP, ISL-MP, or TDF (Table S3). We then observed the extent of excision and rescue without ATP or PP_i_ to assess if rescue was detectable in unfavorable primer unblocking conditions. We saw a rescue deficiency across ddAMP and TDF conditions (Fig. S7B), though ISL-MP-terminated DNA could still be extended by RT_WT_ and RT_M184V_ in 30.5% and 19.6% of respective complexes (Table S5A). These two averages significantly differed from other RTs from BH-corrected t-tests at *Q* = 0.01.

We introduced ATP as an excision-mediating cofactor, yet we noticed minimal DNA rescue with ddAMP or TDF as the primer-terminating base (Fig. 5B). Contrastingly, ISL-MP-terminated DNA was extended to differing levels among RT variants, with RT_WT_ presenting the highest activity and extending up to ∼50% of substrate (Fig. 5B). Among the single mutants, RT_M184V_ leveled off at ∼25% extension, and RT_A114S_ reached just above 10% rescue by 90 mins (Fig. 5B). RT_M184V/A114S_ showed minimal DNA rescue over the 90-min period (Fig. 5B).

We then tested PP_i_ as the excision-mediating cofactor. Interestingly, TDF appeared most prone to PP_i_-mediated excision, with near-identical, rapid excision by all RT variants (Fig. 5C). RT_WT_ and RT_M184V_ efficiently rescued ddAMP-terminated primer while RT_A114S_ and RT_M184V/A114S_ required the full 90 mins to achieve such levels (Fig. 5C); nonetheless, all four variants rescued ∼80% of DNA (Fig. 5C). Excluding RT_M184V/A114S_, the RT variants comparably rescued the full ISL-MP-terminated primer population efficiently, with RT_A114S_ showing a minor lag (Fig. 5C). The double-mutant could not recapitulate these rescue levels, reaching only ∼60% rescue (Fig. 5C). Nonetheless, with no complete deficiencies in excision across conditions, we conclude that excision is not a major factor in ISL resistance or TDF hypersusceptibility.

### Combination experiments reveal additive effects of using ISL and TDF to treat HIV-1

We next tested *in cellulo* antiviral potencies against the four HIV-1 variants-of-interest: WT, M184V/A114S, M184V, and A114S. We treated TZM-GFP cells with ISL or TDF and then conducted single-round infection using each VSV-G pseudotyped HIV-1 variant. We equated GFP puncta to infected cell count. Inhibition curves revealed ISL resistance and TDF hypersusceptibility in pseudotyped virus mutants (Figs. 6A, S12A). Targeting the double mutant produced the largest fold-changes in half-maximal inhibitory concentrations (IC_50_s), where pseudotyped HIV-1_M184V/A114S_ demonstrated 19.3-fold ISL resistance and 10.1-fold TDF hypersusceptibility *vs.* WT (Fig. S12B-C).

**Fig. 6.**
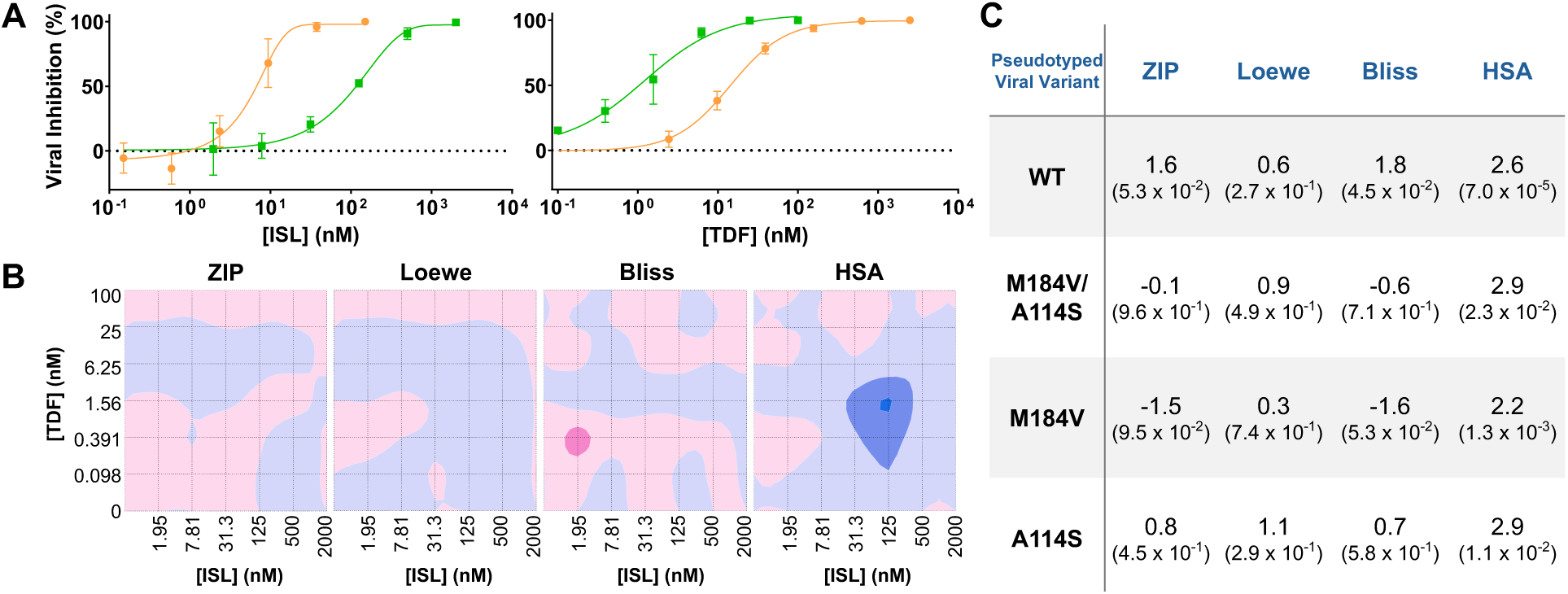
*In cellulo* pseudotyped viral inhibition and combination profiles of ISL and TDF. *[A]* Results of treating of VSV-G pseudotyped WT (orange) and M184V/A114S (green) HIV-1 with ISL and TDF. Values reported from results of three experiments. Graphs made in GraphPad Prism 9, with calculations based on four-parameter inhibition. *[B]* ISL/TDF synergy landscapes resulting from inhibition of pseudotyped M184V/A114S virus across the ZIP, Loewe, Bliss, and HSA models. Synergy Scores and maps were calculated using SynergyFinder+. *[C]* Average Synergy Scores of ISL + TDF (+10 ≥ Mean Synergy Score ≥ -10). Synergy maps plotted for -30 ≥ Synergy Score ≥ +30. Corresponding p-value for each Synergy Score reported within parentheses.

Given these contrasting profiles, we tested the synergistic potential of ISL and TDF against the VSV-G pseudotyped HIV-1 variants. We obtained maps displaying a Synergy Score at each dosing combination based on viral inhibition. From the Zero Interaction Potency (ZIP), Loewe, Bliss, and Highest Single Agent (HSA) synergy models, all 16 landscapes depicted regions of higher and lower scores (Figs. 6B, S13), and the average Synergy Score of each landscape revealed that ISL and TDF work additively (+10 ≥ mean ≥ -10) to target all HIV-1 variants-of-interest (Fig. 6C). Thus, these results provide support for the design of treatment strategies using ISL and TDF to target WT and ISL-resistant HIV-1 variants.

## DISCUSSION

Because HIV-1 RT is the most-commonly targeted viral enzyme by ARTs, resistant mutants commonly appear upon treatment. Drug-resistant infections warrant the need for novel ART regimens. Historically, M184V has been a well-characterized mutation-of-concern, posing clinical resistance or sensitization to select NRTIs^24, 25^. Therefore, it is unsurprising that NRTTI treatment would select for M184V as well. The addition of A114S to M184V heightens concerns over resistance, as the role of A114S in NRTI resistance has not been described^35, 36^. Though studies have identified TDF hypersusceptibility in VSV-G pseudotyped HIV-1_M184V/A114S and M184V_^29^, there has been little follow-up on the mechanism of this hypersensitization. In addition to cell passaging^29^, ISL and TDF have been suggested as a compatible, post-first-line treatment for multidrug-resistant cases^37^. Based on our results, we propose ISL and TDF as a novel treatment strategy against drug-resistant HIV-1 variants.

Before comparing our structures harboring M184V and/or A114S, we proposed that steric crowding at the polymerase active site would cause ISL-TP resistance. This was informed by *[1]* our consideration of residues, like M184, that mutate to β-branched amino acids to confer steric hindrance and *[2]* the flanking nature of residues 114 and 184 around the inhibitor^27^. From our double mutant structure (PDB ID 12SK), V184 appeared closer to the 4’-E of ISL-TP than M184, and S114 had multiple close-range contacts with Mg^2+^-coordinated residues to stabilize their positions (Fig. 1B-D). We believe this combination prevents ISL-TP from evading V184 hindrance. Thus, our structures overall supported ISL-TP resistance occurring during inhibitor incorporation^38^.

When comparing *in cellulo* and *in vitro* inhibitor activities, we noted patterns of TDF hypersusceptibility and ISL resistance among mutants. The largest changes to antiviral efficacies occurred with both mutations present, causing 19-fold ISL resistance in pseudotyped HIV_M184V/A114S_ and 116-fold resistance during *in vitro* primer extension by RT_M184V/A114S_ (Figs. 3, 6). During single-step incorporation, there was a 308-fold decrease in ISL-TP incorporation efficiency by RT_M184V/A114S_ *vs.* RT_WT_ due to the 239-fold *K*_m_ reduction (Table 1). The lower ISL-TP affinity of RT_M184V/A114S_ comes from the combination of both mutations, as the single mutants each possessed only slightly lessened ISL-TP affinity (Fig. 4). The high *in vitro* resistance is consistent with previously reported >100-fold resistance during pre-steady state incorporation kinetics and prior suggestions that ISL-TP binding affinity is affected by M184V and A114S^22, 30^. In agreement with our structural observations of active site steric crowding, RT_M184V/A114S_ confers ISL-TP resistance by inefficiently incorporating the inhibitor (Fig. 1, Table 1).

Our TDF-DP structures revealed spatial interaction benefits conferred by both mutations. In the double mutant structure (PDB ID 12SW), V184 eased steric crowding near the methoxypropyl component of the molecule while S114 participated in an H-bonding interaction with the TDF-DP β-phosphate, suggesting an increase in favorable antiviral-protein interactions (Fig. 2B-C). We also noted that TDF-DP has been shown to undergo non-canonical Watson-Crick (WC) base pairing with dTMP and assessed this in our structures^31^. We instead saw that the density of the TDF-DP/dTMP H-bonding interface looked like that of WC-paired dAMP/dTMP in the same chain. Overall, our structures suggested TDF-DP hypersusceptibility occurs during incorporation.

Interestingly, we did not observe changes to TDF-DP incorporation kinetics with RT_M184V/A114S_ (Table 1, Fig. 4). Though the *K*_m_ of both dATP and TDF-DP worsened with the double mutant, the TDF-DP SI increased by 30% (Table 1), indicating the ratio of TDF-DP to dATP incorporation efficiency is improved in RT_M184V/A114S_. To determine if the improved SI resulted from TDF-DP acting as a better substrate and/or dATP acting as a worse substrate for the double mutant, we calculated the TDF-DP inhibitory constant (*K*_i_). Since inhibitor kinetics were not tested in the presence of competitive substrate, this estimate would consider inhibitor affinity in the presence of dATP. We observed a *K*_i_ improvement for TDF-DP with the double mutant (Table 1), suggesting dATP acts as a worse substrate for RT_M184V/A114S_. Therefore, we posit that the worsened dATP kinetics – captured by *K*_i_ but not steady-state incorporation kinetics alone – cause TDF-DP hypersusceptibility during incorporation (Table 1).

When incorporated into a nascent strand, the inhibitor sits in the *N* site, but RT translocation to the *P* complex allows an incoming base to enter the emptied *N* site (Fig. S1B). This would form a fixed, inhibited system termed a “dead-end complex” (DEC)^39, 40^, yet RT can utilize ATP or PP_i_ to excise the *N* site inhibitor before the fatal DEC is formed^41^. Our translocation and inhibitor excision experiments did not show any patterns of DEC formation, as all RTs substantially translocated and performed successful PP_i_-mediated excision (Fig. 5A). Consistent with previous reports, TDF could readily translocate even at low levels of incoming dTTP (Fig. 5A) ^42^. TDF can be facilely excised by PP_i_ (Fig. 5C)^42^, but RT_M184V/A114S_ inefficiently translocated along TDF-terminated DNA (Fig. 5A). This appeared counterintuitive for a hypersensitive mechanism: if TDF-DP remains at the excision-accessible *N* site, does this suggest TDF-DP resistance at this step? We propose some explanations: *[1]* translocation at physiological dTTP levels is not variable enough among RT variants to justify significant differences, *[2]* inefficient translocation in high dTTP shows that RT_M184V/A114S_ is locked in an unproductive position, or *[3]* as reported before, TDF-DP can adopt an alternate conformation that hinders RT activity^31^. Additional studies are needed to fully understand these observations.

While evaluating inhibitor excision, our experiments examined enzyme fitness (rescue). Effectively, we assessed the paired excision and incorporation activities of each RT variant, the latter which could also be interpolated from steady-state incorporation assays. It is crucial to consider, then, that excision can be inefficiently catalyzed by ATP or dATP^43^. This justifies the slight increase in baseline rescue across controls (Fig. S7B), as d(d)NTP rescue mixes contained 100 µM dATP to outcompete inhibitors. Still, it is well-known that ATP-mediated excision activity is low without thymidine analog resistance mutations (TAMs)^43^, which are not studied here. Hence, we believe the apparent “ATP-mediated excision and rescue” in the case of ISL-MP instead was purely rescue; d(d)NTPs could be incorporated through the chemically competent 3’-OH of ISL-MP without prior excision of the inhibitor (Fig. 5B). In support of this, rescue levels among the RT variants followed the pattern of the ATP/PP_i_-absent controls and mirrored the fitness levels in dATP incorporation tests (Table 1, Fig. S7B). Thus, we believe any differences in this assay stem from inherent differences during incorporation, where both ISL resistance and TDF hypersusceptibility occur.

Observing resistance and hypersusceptibility to two antivirals is not unique to M184V and A114S, with complementary pairings arising in other nucleoside and/or non-nucleoside analog combinations as well^44, 45^. Nonetheless, we explored this unique instance in which the mutations respond contrastingly to analogs of the same nucleotide: dATP. The ability of ISL and TDF to work harmoniously stems from rate-limiting inhibitor phosphorylation^20, 46^. Instead of competing for the same kinase, ISL becomes ISL-TP using deoxycytidine kinase (DCK), and TDF becomes TDF-DP, first by adenylate kinase 2 and then pyruvate and creatine kinases^47, 48^. Correspondingly, an additive effect from combining ISL and TDF had been predicted based on the MacSynergy II method^49^, which our cell-based combination experiments across four models confirmed (Figs. 6, S13).

There are potential limitations of this study. Firstly, we qualify that structural assertions were made within the resolution constraints of our structures. Nonetheless, to our knowledge, our RT_M184V_/dsDNA/TDF-DP structure is the highest-resolution structure to-date of HIV-1 RT targeted by TDF-DP, determined to 2.63 Å. Moreover, we conducted *in cellulo* studies using subtype B-derived, pseudotyped HIV-1 particles and acknowledge that this does not represent the full breadth of circulating HIV strains. In-tandem, we did not perform *in vitro* competition assays that would corroborate *in cellulo* ISL/TDF additivity. Nevertheless, we contend that whether TDF or ISL binds to the DNA primer, both can potently inhibit RT; prior work has also shown that each can have preferred, sequence-specific binding locations, together expanding the available targets on the DNA strand compared to either antiviral alone^10, 11^. Moreover, we believe that in using longer templates for *in vitro* primer extension, inhibitor activities would be more defined. Taking into account the full viral genome, the complementary antiviral profiles presented here would be further evident. Also, there were additional bands visible in both translocation and excision gels (Figs. S10, S11). In footprinting gels, there were bands upstream and downstream of expected bands (Fig. S10); we ascribe this to non-specific cleavage by hydroxyl radicals, as these bands appeared in our ddAMP and ISL-MP gels and in previous studies^10^. In excision gels of the ddAMP-and TDF-terminated DNA, bands appeared below the P_1_ inhibitor-terminated band (Fig. S11). We hypothesize that there is a primer subset where excision occurred, yet primer was not rescued due to enzyme fitness defects.

Considering ISL has just received FDA approval as the first-in-class NRTTI, preemptive characterization of ARTs circumventing drug resistance is needed. Ongoing antiviral development efforts have resulted in MK-8527 as a next-generation NRTTI inspired by ISL^50^. While in the current study, we evaluate ISL resistance mutations that present TDF hypersusceptibility, in previous studies, we have also highlighted TDF resistance mutations such as the K65R mutation that conversely presents ISL hypersusceptibility^51^. This scenario further supports ISL/TDF for the same reason: one antiviral’s weakness may be the other’s strength. Hence, we emphasize the utility of taking inspiration from these two compounds toward new ART strategies that can treat ART-resistant HIV-1 infections.

## METHODS

### VIROLOGICAL STUDIES

All virological assays were performed and analyzed as previously described^52^. Any modifications are provided below. Results were analyzed and graphed in GraphPad Prism 9.

### Assessment of synergistic potential of combined ISL + TDF treatment

The NEBuilder® HiFi DNA Assembly Kit (New England Biolabs) was used to introduce the M184V, A114S, and M184V/A114S mutations into the HIV-1 NL4-3 *Δenv* construct. Whole-plasmid sequencing (AZENTA Genewiz) confirmed presence of each mutation and preservation of the HIV-1 NL4-3 *Δenv* backbone. The TDF prodrug (NIH HIV Reagent Program) was used.

| Effect of Combined Treatment | Average Synergy Score (x) |
| --- | --- |
| Synergistic | $x \geq +10$ |
| Additive | $+10 > x > -10$ |
| Antagonistic | $x \leq -10$ |

All biochemical assays were performed highly similar to previously described methods^53^, with any additional modifications listed below. Results were analyzed and graphed in GraphPad Prism 9.

### Preparation of biological reagents for *in vitro* mechanistic assays

Through the Emory Integrated Genomics Core (EIGC), the M184V and/or A114S mutations were cloned into the biochemistry-relevant HIV-1 RT p66 construct^10, 11^. HIV-1 RT p66 plasmids were co-transformed with a p51-encoding plasmid into JM109 competent *E. coli* cells (Agilent Technologies, Inc.). Cells were grown in LB with 1X streptomycin, 1X kanamycin, and 1X chloramphenicol at 37°C, and upon reaching an OD_600_ of ∼0.7-0.9, RT expression was induced with 1.67 mM of isopropyl 𝛽-D-1-thiogalactopyranoside (IPTG; Fisher Scientific Company, LLC.) and left for 3 h. After centrifugation, resuspended pellets were treated with lysozyme, nucleases, sonication, and centrifugation at 18,000 g for 30 m at 4°C. RTs were then FPLC-purified from 0.45 µm filter-clarified lysates using tandem nickel-affinity (Cytiva) and heparin-affinity (Cytiva) chromatography. Fractions containing RT were pooled, concentrated, and stored at -80°C. Protein yield and purity were confirmed by SDS-PAGE using Novex™ 4-12% Polyacrylamide Tris-Glycine Mini Protein Gels (Novex Products, Inc.).

A Cyanine3 (Cy3) fluorophore was placed on the 5’-termini of select strands (Table S3). All dsDNA substrates were formed by mixing a 3:1 molar ratio of template to primer, denaturing at 95°C for 5 m, and reannealing at room temperature for 30 m. Stocks were stored at -20°C in darkened conditions.

### *In vitro* antiviral susceptibility against primer extension activity

10X stocks of antiviral – islatravir triphosphate triethylamine salt (ISL-TP TEA; TargetMol Chemicals, Inc.) or tenofovir diphosphate triethylamine salt (TDF-DP TEA; TargetMol Chemicals, Inc.) – were prepared in nuclease-free water, ranging from 0 to 50 µM.

### Steady state nucleotide incorporation

**i. Time dependence**

Reaction mixes were incubated at 37°C, from which timepoints were quenched in an equal volume of 100% formamide up to 60 m.

**ii. Concentration dependence**

Based on densitometry results, the 3-m timepoint was determined to ubiquitously fall within the linear range of incorporation rates across all 12 RT/nucleotide combinations. 10X stocks of dATP (Thermo Scientific™) or inhibitor were prepared in nuclease-free H_2_O, ranging from 0 to 250 µM. Reactions were incubated at 37°C for 3 m, quenched in an equal volume of 100% formamide, and heat shocked for 5 m at 95°C.

### Hydroxyl-radical DNA footprinting to assess HIV-1 RT translocation state

This assay was performed based on what has been previously described^53^.

### ATP-/PP_i_-dependent nucleotide excision and rescue of inhibitor-terminated primers

**i. Preparation of pyrophosphatase-treated nucleotides and inhibitor-terminated DNA substrates**

Inhibitors (ddATP, ISL-TP, TDF-DP) and nucleotides (ATP, dATP, dTTP, ddGTP [AAT Bioquest, Inc.]) were pyrophosphatase treated in master-mixes (1 mM DTT, 50 mM Tris-HCl pH 8.0, 50 mM KCl, 60 mM MgCl_2_, 0.5 units pyrophosphatase [Sigma-Aldrich Co.]). Reactions were incubated for 1 h at 37°C prior to centrifugation at 13,000 g for 30 m at 4°C, and a Nanodrop was used to determine nucleotide concentration. Treated inhibitors were used to terminate P_d18_-P_0_-Cy3 annealed to T_d31_ in a 1:3 molar ratio.

**ii. ATP-/PPi-based excision and rescue reactions**

This assay was performed based on what has been previously described^53^.

### Denaturing gel densitometry

This assay was performed based on what has been previously described^54^.

## STRUCTURAL STUDIES

### Preparation of ternary complexes for structural studies

Through the EIGC, the M184V and/or A114S mutations were cloned into the previously characterized pCDFDuet-1 vector containing the WT p66 subunit with the Q258C mutation for primer crosslinking^55^. HIV-1 RT p66 plasmid was co-transformed with pRSFDuet-1 backbone containing the p51 subunit into BL21-CodonPlus (DE3)-RIL competent cells (Agilent Technologies, Inc.). Cells were grown in LB at 37°C, at an OD_600_ of ∼0.7-0.9, RT expression was induced with 1.67 mM of IPTG for 3 h. After centrifugation, resuspended pellets were treated with lysozyme, nucleases, sonication, and centrifugation. RTs were FPLC-purified using nickel-affinity (Cytiva) followed by Heparin-affinity (Cytiva) chromatography^12^. Fractions containing RT were concentrated and stored at -80°C in 10 mM Tris-HCl pH 8.0 and 75 mM NaCl. Protein yield and purity were confirmed by SDS-PAGE at each purification step using Novex™ 4-12% Polyacrylamide Tris-Glycine Mini Protein Gels (Novex Products, Inc.).

Overnight protein-DNA crosslinking was conducted at 30°C in 25 mM Tris-HCl pH 8.0, 100 mM NaCl, 15 µM P_d18_*/T_d24_, 100 µM ddGTP, 2 mg RT, and 10 mM MgCl_2_^12^. RT-DNA binary complexes were purified using tandem nickel-affinity (Cytiva) and Heparin affinity (Cytiva) chromatography and concentrated to 12-14 mg/mL.

### X-ray crystallography

Crosslinked RT-DNA (at 10 mg/mL RT) was incubated with 5 mM MgCl_2_ and 1.5X molar excess ISL-TP or 10X molar excess TDF-DP at room temperature for 1 h. Hanging-drop vapor diffusion was conducted in 24-well pre-greased plates (Greiner Bio-One International GmbH) containing 500-µL of well solution and 1- to 3-µL drops in 2:1, 1:1, and 1:2 protein-to-solution ratios. Crystals grew in 25 mM Bis-Tris Propane pH 5.8-6.2 (AmBeed, Inc.); 5-10% (w/v) PEG 4,000 or 8,000 (Hampton Research Corp.); and 5 mM MgCl_2_. Crystals were cryoprotected in 20-24% ethylene glycol (Hampton Research Corp.) up to 1 m and cryo-cooled in LN_2_.

Crustal pucks were dry-shipped to Brookhaven National Laboratory (BNL) National Synchrotron Light Source II AMX Beamline or Advanced Light Source 4.2.2 Beamline for data collection on a Dectris Eiger 9M detector or an RDI CMOS-8M Detector, respectively^56, 57^.

All datasets were processed using XDS, indexed in space group C 2 2 2_1_ and with one HIV-1 RT molecule in the asymmetric unit^58^. Each dataset was analyzed in XTRIAGE to confirm there was no twinning present^59^. Structure resolutions and unit cell dimensions for each crystal structure are provided below:

| Complex-of-Interest | Resolution | Unit Cell Dimensions (a, b, c) |
| --- | --- | --- |
| RT<sub>M184V/A114S</sub>/dsDNA<sub>ddGMP</sub>/ISL-TP | 2.79 Å | 168.97Å, 171.08Å, 106.27Å |
| RT<sub>M184V</sub>/dsDNA<sub>ddGMP</sub>/ISL-TP | 2.91 Å | 167.59Å, 169.50Å, 101.68Å |
| RT<sub>A114S</sub>/dsDNA<sub>ddGMP</sub>/ISL-TP | 2.65 Å | 167.52Å, 170.56Å, 105.31Å |
| RT<sub>M184V/A114S</sub>/dsDNA<sub>ddGMP</sub>/TDF-DP | 3.20 Å | 165.71Å, 172.25Å, 105.42Å |

Initial phases were solved by molecular replacement using Phaser, using as a starting model the coordinates of WT HIV-1 RT in-complex with dsDNA and ISL-TP (PDB ID 5J2M) with all ligands and solvent molecules removed^12, 60^.

### Cryogenic electron microscopy

Crosslinked RT-DNA (at 80 µM RT) was incubated with 12.5X molar excess TDF-DP in 10 mM Tris-HCl pH 8.0, 250 mM NaCl, 10 mM MgCl_2_, and 0.2% [w/v] n-Octyl-β-D-glucopyranoside (Thermo Scientific Chemicals, Inc.) at room temperature for 1 h. UltraAufoil® R 1.2/1.3 Holey Gold 300-Mesh Film grids (Quantifoil Micro Tools GmbH) were glow-discharged using a GloQube® Plus (Quorum; 15 s at 20 mA) and plunge-frozen into LN_2_ (4 µL sample, 1.5-s blot time, 10-s wait time, 0% blot force, 100% humidity) using the Vitrobot Mark IV System (ThermoFisher). Grids were prepared through the Emory Robert P. Apkarian Integrated Electron Microscopy Core (IEMC).

**i. RT_M184V_/dsDNA/TDF-DP**: Grids were screened using the IEMC’s Thermo Scientific™ Talos Arctica™ Cryo-TEM prior to shipment to the BNL Laboratory for BioMolecular Structure for collection on a Thermo Scientific™ Krios™ G3i (300 kV) Cryo-TEM using a Gatan K3 camera. Datasets were processed in CryoSPARC v4.7.1 to 2.87 Å.
**ii. RT_A114S_/dsDNA/TDF-DP**: Grids were screened and collected on at 100kX magnification using the IEMC’s Thermo Scientific™ Talos Arctica™ (200 kV) Cryo-TEM using a Gatan K3 camera and BioQuantum energy filter. Datasets were processed in CryoSPARC v4.7.1 to 2.63 Å.

All six structures were iteratively refined using Phenix v2.0-5885 and Coot v0.9.8.95^61, 62^. Final models were validated using MolProbity and the PDB Validation Server^63^. Some of the structural biology applications used in this project were compiled and configured by SBGrid^64^. Maps and coordinates were deposited into the wwPDB for public access^65^. All PDB accession codes, data collection information, processing workflows, and refinement statistics are available in Supplementary Tables S1-2.

## ADDITIONAL FORMULAE

### i. Inhibitory constant (*K*_i_)

The Cheng-Prusoff equation for a competitive inhibitor under non-cooperative assumptions was used:

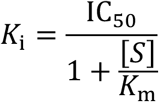

*IC_50_: half-maximal* in vitro *inhibitory concentration; S: 10* 𝜇*M dATP; K*_m_*: dATP Michaelis constant*

### ii. Propagation of error

Fold-changes in *in vitro* and *in vivo* responses to antivirals required propagation of the error reported for each sample involved in fold-change calculations. The standard error for a given sample was first calculated from its standard deviation using the following formula:

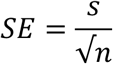

*SE: standard error; s: standard deviation; n: sample size*

The “propagation of standard error” formula for the quotient of independent values was used to calculate catalytic efficiency (*k*_cat_/*K*_m_) and fold-change as follows, with variables defined below:

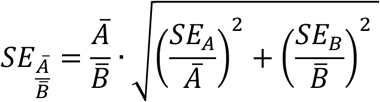

*A: first sample-of-interest; B: second sample-of-interest; SE_X_: standard error of given sample; x̄: average of given sample*

## ACKNOWLEDGEMENTS AND CONTRIBUTIONS

We thank the fellow members of the Sarafianos Laboratory for their plentiful insights. We also acknowledge Dr. Zhe Li Salie and Dr. Bruno Marchand for their extensive contributions to crystallographic studies, along with Dr. Srihari Nagendra Ravi Koripella and Dr. Robert Dick for their guidance toward cryo-EM studies. S.M.R. thanks the Cold Spring Harbor Laboratory Macromolecular Crystallography Course and the National Center for Cryo-EM Access and Training for providing invaluable training.

These studies were supported in-part by the Emory Integrated Genomics Core (EIGC; RRID:SCR_023529), which is subsidized by the Emory University School of Medicine and is one of the Emory Integrated Core Facilities. Additional support was provided by the Georgia Clinical & Translational Science Alliance of the National Institutes of Health under Award Number UL1TR002378. This work was additionally supported by the Emory University Robert P. Apkarian Integrated Electron Microscopy Core Facility (RRID: SCR_023537), which is subsidized by the School of Medicine and Emory College of Arts and Sciences. Additional support was provided by the Georgia Clinical & Translational Science Alliance of the National Institutes of Health under award number UL1TR000454.

This research used resources of the Advanced Light Source, which is a DOE Office of Science User Facility under contract no. DE-AC02-05CH11231. Beamline 4.2.2 of the Advanced Light Source, a DOE Office of Science User Facility under Contract No. DE-AC02-05CH11231, is supported in part by the ALS-ENABLE program funded by the National Institutes of Health, National Institute of General Medical Sciences, Grant No. P30GM124169-01. This research also used resource 17-ID-1 of the National Synchrotron Light Source II, a U.S. Department of Energy Office of Science User Facility operated for the DOE Office of Science by Brookhaven National Laboratory under Contract No. DE-SC0012704. The Center for BioMolecular Structure is primarily supported by the National Institutes of Health, National Institute of General Medical Sciences through Grant No. P30GM133893, and by the DOE Office of Biological and Environmental Research FWP No. BO070. This research was additionally supported in-part by the Brookhaven National Laboratory (BNL) Laboratory for BioMolecular Structure (LBMS). LBMS is supported by the DOE Office of Biological and Environmental Research (KP1607011). Some of the structural biology applications used in this project were compiled and configured by SBGrid.

This work was supported in part by the National Institutes of Health (Grants R37AI076119 and P30AI050409 to S.G.S.; F31AI194923 to S.M.R.; F31AI172618 to A.A.S.; S.M.R. and A.A.S. were supported in-part by T32GM135060). The content is solely the responsibility of the authors and does not necessarily reflect the official views of the National Institutes of Health. S.G.S. acknowledges the Nahmias-Schinazi Distinguished Chair in Research. S.M.R. acknowledges support from the Robert W. Woodruff Foundation.

## Author Contributions

Conceptualization: S.M.R., A.A.S., E.M., and S.G.S.; Methodology & Investigation: S.M.R., A.A.S., K.H.K., I.L.K., X.W., E.M., and K.A.K.; Writing – Original Draft: S.M.R.; Writing – Review & Editing: S.M.R., A.A.S., K.A.K., and S.G.S.

## REFERENCES

1. Research Toward HIV Cure: National Institutes of Health | Office of AIDS Research; 2025. Available from: https://www.oar.nih.gov/hiv-policy-and-research/research-priorities-overview/research-toward-hiv-cure.

2. Gaebler C. The Next Berlin patient: Sustained HIV Remission Surpassing Five Years Without Antiretroviral Therapy After Heterozygous CCR5 WT/Δ32 Allogeneic Hematopoietic Stem Cell Transplantation. 25th International AIDS Conference: Charité - Universitätsmedizin Berlin, 2024 July 22-26, 2024. Report No.

3. Xiao Q, He S, Wang C, Zhou Y, Zeng C, Liu J, Liu T, Li T, Quan X, Wang L, Zhai L, Liu Y, Li J, Zhang X, Liu Y. Deep Thought on the HIV Cured Cases: Where Have We Been and What Lies Ahead? Biomolecules. 2025;15(3):378. doi: 10.3390/biom15030378.

4. Gandhi RT, Landovitz RJ, Sax PE, Smith DM, Springer SA, Günthard HF, Thompson MA, Bedimo RJ, Benson CA, Buchbinder SP, Crabtree-Ramirez BE, Del Rio C, Eaton EF, Eron JJ, Hoy JF, Lehmann C, Molina J-M, Jacobsen DM, Saag MS. Antiretroviral Drugs for Treatment and Prevention of HIV in Adults: 2024 Recommendations of the International Antiviral Society–USA Panel. JAMA. 2025;333(7):609. doi: 10.1001/jama.2024.24543.

5. Ravichandran SM, McFadden WM, Snyder AA, Sarafianos SG. State of the ART (antiretroviral therapy): Long-acting HIV-1 therapeutics. Global Health & Medicine. 2024;6(5):285–94. doi: 10.35772/ghm.2024.01049.

6. HIVdb. HIV Drug Resistance Database: Stanford University; 2026. Available from: https://hivdb.stanford.edu/.

7. Tao K, Zhou J, Nagarajan P, Tzou PL, Shafer RW. Comprehensive Database of HIV mutations Selected During Antiretroviral In Vitro Passage Experiments. Antiviral Research. 2024;230:105988. doi: 10.1016/j.antiviral.2024.105988.

8. HIVinfo. FDA-Approved HIV Medicines HIVinfo.NIH.gov2025. Available from: https://hivinfo.nih.gov/understanding-hiv/fact-sheets/fda-approved-hiv-medicines.

9. Singh AK, Das K. Insights into HIV-1 Reverse Transcriptase (RT) Inhibition and Drug Resistance from Thirty Years of Structural Studies. Viruses. 2022;14(5):1027. doi: 10.3390/v14051027.

10. Michailidis E, Marchand B, Kodama EN, Singh K, Matsuoka M, Kirby KA, Ryan EM, Sawani AM, Nagy E, Ashida N, Mitsuya H, Parniak MA, Sarafianos SG. Mechanism of Inhibition of HIV-1 Reverse Transcriptase by 4′-Ethynyl-2-fluoro-2′-deoxyadenosine Triphosphate, a Translocation-defective Reverse Transcriptase Inhibitor. Journal of Biological Chemistry. 2009;284(51):35681–91. doi: 10.1074/jbc.m109.036616.

11. Michailidis E, Huber AD, Ryan EM, Ong YT, Leslie MD, Matzek KB, Singh K, Marchand B, Hagedorn AN, Kirby KA, Rohan LC, Kodama EN, Mitsuya H, Parniak MA, Sarafianos SG. 4′-Ethynyl-2-fluoro-2′-deoxyadenosine (EFdA) Inhibits HIV-1 Reverse Transcriptase with Multiple Mechanisms. Journal of Biological Chemistry. 2014;289(35):24533–48. doi: 10.1074/jbc.m114.562694.

12. Salie ZL, Kirby KA, Michailidis E, Marchand B, Singh K, Rohan LC, Kodama EN, Mitsuya H, Parniak MA, Sarafianos SG. Structural Basis of HIV Inhibition by Translocation-defective RT Inhibitor 4′-ethynyl-2-fluoro-2′-deoxyadenosine (EFdA). Proceedings of the National Academy of Sciences. 2016;113(33):9274–9. doi: 10.1073/pnas.1605223113.

13. Rockstroh JK, Paredes R, Cahn P, Molina J-M, Sokhela SM, Hinestrosa F, Kassim S, Cunningham D, Ghosn J, Bogner JR, Gatanaga H, Asante-Appiah E, Zhang Y, Nwoke U, Klopfer SO, Eves K, Squires K, Correll T, Fox MC, Pisculli ML. Doravirine/Islatravir (100/0.75 mg) Once-Daily Compared With Bictegravir/Emtricitabine/Tenofovir Alafenamide as Initial HIV-1 Treatment: 48-Week Results From a Phase 3, Randomized, Controlled, Double-Blind, Noninferiority Trial. Clinical Infectious Diseases. 2025;81(2):322–32. doi: 10.1093/cid/ciaf077.

14. Vanderveen LA, Selzer L, Chang S, Li J, Diamond TL, Asante-Appiah E, Llamoso C, Rhee MS, Callebaut C. Resistance Analysis of Weekly Islatravir Plus Lenacapavir in People with HIV at 48 Weeks. JAIDS Journal of Acquired Immune Deficiency Syndromes. 2026. doi: 10.1097/qai.0000000000003873.

15. Sension MG, Brunet L, Hsu RK, Fusco JS, Cochran Q, Uranaka C, Sridhar G, Vannappagari V, Van Wyk J, McCurdy L, Wohlfeiler MB, Fusco GP. Cabotegravir + Rilpivirine Long-Acting Injections for HIV Treatment in the US: Real World Data from the OPERA Cohort. Infectious Diseases and Therapy. 2023;12(12):2807–17. doi: 10.1007/s40121-023-00890-2.

16. Wang H, Lu X, Yang X, Xu N. The efficacy and safety of tenofovir alafenamide versus tenofovir disoproxil fumarate in antiretroviral regimens for HIV-1 therapy. Medicine. 2016;95(41):e5146. doi: 10.1097/md.0000000000005146.

17. Osuala EC, Naidoo A, Dooley KE, Naidoo K, Perumal R. Broadening access to tenofovir alafenamide for the treatment and prevention of HIV-1 infection. Expert Review of Clinical Pharmacology. 2023;16(10):939–57. doi: 10.1080/17512433.2023.2251387.

18. Lovett GC, Nguyen T, Iser DM, Holmes JA, Chen R, Demediuk B, Shaw G, Bell SJ, Desmond PV, Thompson AJ. Efficacy and safety of tenofovir in chronic hepatitis B: Australian real world experience. World Journal of Hepatology. 2017;9(1):48. doi: 10.4254/wjh.v9.i1.48.

19. Chan HLY, Buti M, Lim Y-S, Agarwal K, Marcellin P, Brunetto M, Chuang W-L, Janssen HLA, Fung S, Izumi N, Abdurakhmanov D, Jabłkowski M, Celen MK, Ma X, Caruntu F, Flaherty JF, Abramov F, Wang H, Camus G, Osinusi A, Pan CQ, Shalimar, Seto W-K, Gane E. Long-Term Treatment With Tenofovir Alafenamide for Chronic Hepatitis B Results in High Rates of Viral Suppression and Favorable Renal and Bone Safety. American Journal of Gastroenterology. 2024;119(3):486–96. doi: 10.14309/ajg.0000000000002468.

20. Holec AD, Mandal S, Prathipati PK, Destache CJ. Nucleotide Reverse Transcriptase Inhibitors: A Thorough Review, Present Status and Future Perspective as HIV Therapeutics. Current HIV Research. 2018;15(6). doi: 10.2174/1570162x15666171120110145.

21. Merck & Co. I. FDA Approves Merck’s Once-Daily IDVYNSO™ (doravirine/islatravir). 2026.

22. Cilento ME, Reeve AB, Michailidis E, Ilina TV, Nagy E, Mitsuya H, Parniak MA, Tedbury PR, Sarafianos SG. Development of Human Immunodeficiency Virus Type 1 Resistance to 4′-Ethynyl-2-Fluoro-2′-Deoxyadenosine Starting with Wild-Type or Nucleoside Reverse Transcriptase Inhibitor-Resistant Strains. Antimicrobial Agents and Chemotherapy. 2021;65(12). doi: 10.1128/aac.01167-21.

23. Diamond TL, Ngo W, Xu M, Goh SL, Rodriguez S, Lai M-T, Asante-Appiah E, Grobler JA. Islatravir Has a High Barrier to Resistance and Exhibits a Differentiated Resistance Profile from Approved Nucleoside Reverse Transcriptase Inhibitors (NRTIs). Antimicrobial Agents and Chemotherapy. 2022;66(6). doi: 10.1128/aac.00133-22.

24. GöTte M, Arion D, Parniak MA, Wainberg MA. The M184V Mutation in the Reverse Transcriptase of Human Immunodeficiency Virus Type 1 Impairs Rescue of Chain-Terminated DNA Synthesis. Journal of Virology. 2000;74(8):3579–85. doi: 10.1128/jvi.74.8.3579-3585.2000.

25. Boyer PL, Sarafianos SG, Arnold E, Hughes SH. The M184V Mutation Reduces the Selective Excision of Zidovudine 5′-Monophosphate (AZTMP) by the Reverse Transcriptase of Human Immunodeficiency Virus Type 1. Journal of Virology. 2002;76(7):3248–56. doi: 10.1128/jvi.76.7.3248-3256.2002.

26. Hung M, Tokarsky EJ, Lagpacan L, Zhang L, Suo Z, Lansdon EB. Elucidating molecular interactions of L-nucleotides with HIV-1 reverse transcriptase and mechanism of M184V-caused drug resistance. Communications Biology. 2019;2(1). doi: 10.1038/s42003-019-0706-x.

27. Sarafianos SG, Das K, Clark AD, Ding J, Boyer PL, Hughes SH, Arnold E. Lamivudine (3TC) resistance in HIV-1 reverse transcriptase involves steric hindrance with β-branched amino acids. Proceedings of the National Academy of Sciences. 1999;96(18):10027–32. doi: 10.1073/pnas.96.18.10027.

28. Zang X, Ankrom W, Kraft WK, Vargo R, Stoch SA, Iwamoto M, Matthews RP. Intracellular islatravir-triphosphate half-life supports extended dosing intervals. Antimicrobial Agents and Chemotherapy. 2024;68(9). doi: 10.1128/aac.00458-24.

29. Cilento ME, Wen X, Reeve AB, Ukah OB, Snyder AA, Carrillo CM, Smith CP, Edwards K, Wahoski CC, Kitzler DR, Kodama EN, Mitsuya H, Parniak MA, Tedbury PR, Sarafianos SG. HIV-1 Resistance to Islatravir/Tenofovir Combination Therapy in Wild-Type or NRTI-Resistant Strains of Diverse HIV-1 Subtypes. Viruses. 2023;15(10):1990. doi: 10.3390/v15101990.

30. Zalenski N, Meredith BR, Savoie DJ, Naas MJ, Suo DJ, Betancourt D, Seay TW, Suo Z. Kinetic Investigation of Resistance to Islatravir Conferred by Mutations in HIV-1 Reverse Transcriptase. Journal of Molecular Biology. 2025;437(12):169100. doi: 10.1016/j.jmb.2025.169100.

31. Tuske S, Sarafianos SG, Clark AD, Ding J, Naeger LK, White KL, Miller MD, Gibbs CS, Boyer PL, Clark P, Wang G, Gaffney BL, Jones RA, Jerina DM, Hughes SH, Arnold E. Structures of HIV-1 RT–DNA complexes before and after incorporation of the anti-AIDS drug tenofovir. Nature Structural & Molecular Biology. 2004;11(5):469–74. doi: 10.1038/nsmb760.

32. Cheng H. The influence of cooperativity on the determination of dissociation constants: examination of the Cheng–Prusoff equation, the Scatchard analysis, the Schild analysis and related power equations. Pharmacological Research. 2004;50(1):21–40. doi: 10.1016/j.phrs.2003.11.007.

33. Marchand B, Götte M. Site-specific Footprinting Reveals Differences in the Translocation Status of HIV-1 Reverse Transcriptase. Journal of Biological Chemistry. 2003;278(37):35362–72. doi: 10.1074/jbc.m304262200.

34. Meyer PR, Matsuura SE, So AG, Scott WA. Unblocking of chain-terminated primer by HIV-1 reverse transcriptase through a nucleotide-dependent mechanism. Proceedings of the National Academy of Sciences. 1998;95(23):13471–6. doi: 10.1073/pnas.95.23.13471.

35. Rhee SY. Human immunodeficiency virus reverse transcriptase and protease sequence database. Nucleic Acids Research. 2003;31(1):298–303. doi: 10.1093/nar/gkg100.

36. Rhee S-Y, Kantor R, Katzenstein DA, Camacho R, Morris L, Sirivichayakul S, Jorgensen L, Brigido LF, Schapiro JM, Shafer RW. HIV-1 pol mutation frequency by subtype and treatment experience: extension of the HIVseq program to seven non-B subtypes. AIDS. 2006;20(5):643–51. doi: 10.1097/01.aids.0000216363.36786.2b.

37. Nka AD, Bouba Y, Tsapi Lontsi WR, Gouissi Anguechia D-H, Teto G, Ka’E AC, Semengue ENJ, Ambe Chenwi C, Takou D, Forgwei L, Tekoh TA-K, Ngueko AMK, Fokou BB, Efakika Gabisa J, Tchouaket MCT, Tognapabo WL, Ayuk Ngwese DT, Njiki Bikoi J, Armenia D, Colizzi V, Yotebieng M, Ndembi N, Santoro M-M, Ceccherini-Silberstein F, Perno C-F, Ndjolo A, Fokam J. Tenofovir and Doravirine Are Potential Reverse-Transcriptase Analogs in Combination with the New Reverse-Transcriptase Translocation Inhibitor (Islatravir) Among Treatment-Experienced Patients in Cameroon: Designing Future Treatment Strategies for Low-an. Viruses. 2025;17(1):69. doi: 10.3390/v17010069.

38. Das K, Arnold E. HIV-1 reverse transcriptase and antiviral drug resistance. Part 1. Current Opinion in Virology. 2013;3(2):111-8. doi: 10.1016/j.coviro.2013.03.012.

39. Meyer PR, Matsuura SE, Mian AM, So AG, Scott WA. A Mechanism of AZT Resistance. Molecular Cell. 1999;4(1):35–43. doi: 10.1016/s1097-2765(00)80185-9.

40. Sarafianos SG, Marchand B, Das K, Himmel DM, Parniak MA, Hughes SH, Arnold E. Structure and Function of HIV-1 Reverse Transcriptase: Molecular Mechanisms of Polymerization and Inhibition. Journal of Molecular Biology. 2009;385(3):693–713. doi: 10.1016/j.jmb.2008.10.071.

41. Tu X, Das K, Han Q, Bauman JD, Clark AD, Hou X, Frenkel YV, Gaffney BL, Jones RA, Boyer PL, Hughes SH, Sarafianos SG, Arnold E. Structural basis of HIV-1 resistance to AZT by excision. Nature Structural & Molecular Biology. 2010;17(10):1202–9. doi: 10.1038/nsmb.1908.

42. Marchand B, White KL, Ly JK, Margot NA, Wang R, McDermott M, Miller MD, GöTte M. Effects of the Translocation Status of Human Immunodeficiency Virus Type 1 Reverse Transcriptase on the Efficiency of Excision of Tenofovir. Antimicrobial Agents and Chemotherapy. 2007;51(8):2911–9. doi: 10.1128/aac.00314-07.

43. Acosta-Hoyos AJ, Scott WA. The Role of Nucleotide Excision by Reverse Transcriptase in HIV Drug Resistance. Viruses. 2010;2(2):372–94. doi: 10.3390/v2020372.

44. White KL, Margot NA, Wrin T, Petropoulos CJ, Miller MD, Naeger LK. Molecular Mechanisms of Resistance to Human Immunodeficiency Virus Type 1 with Reverse Transcriptase Mutations K65R and K65R+M184V and Their Effects on Enzyme Function and Viral Replication Capacity. Antimicrobial Agents and Chemotherapy. 2002;46(11):3437–46. doi: 10.1128/aac.46.11.3437-3446.2002.

45. Lai M-T, Feng M, Xu M, Ngo W, Diamond TL, Hwang C, Grobler JA, Hazuda DJ, Asante-Appiah E. Doravirine and Islatravir Have Complementary Resistance Profiles and Create a Combination with a High Barrier to Resistance. Antimicrobial Agents and Chemotherapy. 2022;66(5). doi: 10.1128/aac.02223-21.

46. Mu L, Zhou R, Tang F, Liu X, Li S, Xie F, Xie X, Peng J, Yu P. Intracellular pharmacokinetic study of zidovudine and its phosphorylated metabolites. Acta Pharmaceutica Sinica B. 2016;6(2):158–62. doi: 10.1016/j.apsb.2015.10.002.

47. Kawamoto A, Kodama E, Sarafianos SG, Sakagami Y, Kohgo S, Kitano K, Ashida N, Iwai Y, Hayakawa H, Nakata H, Mitsuya H, Arnold E, Matsuoka M. 2′-Deoxy-4′-C-ethynyl-2-halo-adenosines active against drug-resistant human immunodeficiency virus type 1 variants. The International Journal of Biochemistry & Cell Biology. 2008;40(11):2410–20. doi: 10.1016/j.biocel.2008.04.007.

48. Eberhard CD, Orsburn BC, Bumpus NN. Tenofovir Activation in Brain and Liver is Diminished in Creatine Kinase Brain-Type Knockout Mice. The Journal of Pharmacology and Experimental Therapeutics. 2023;385:383. doi: 10.1124/jpet.122.167260.

49. Hachiya A, Reeve AB, Marchand B, Michailidis E, Ong YT, Kirby KA, Leslie MD, Oka S, Kodama EN, Rohan LC, Mitsuya H, Parniak MA, Sarafianos SG. Evaluation of Combinations of 4′-Ethynyl-2-Fluoro-2′-Deoxyadenosine with Clinically Used Antiretroviral Drugs. Antimicrobial Agents and Chemotherapy. 2013;57(9):4554–8. doi: 10.1128/aac.00283-13.

50. Raheem IT, Girijavallabhan V, Fillgrove KL, Goh SL, Bahnck-Teets C, Huang Q, Li F, Wan B-L, O’Donnell GT, Patteson JB, Cilento ME, Bennet A, Hayes RP, Tummala S, McHale C, Wiltsie J, Ellis J, Asante-Appiah E, Hazuda DJ, Hale J, Grobler JA, Xu M, Diamond TL, Lai M-T. MK-8527 is a novel inhibitor of HIV-1 reverse transcriptase translocation with potential for extended-duration dosing. PLOS Biology. 2025;23(8):e3003308. doi: 10.1371/journal.pbio.3003308.

51. Das K, Bandwar RP, White KL, Feng JY, Sarafianos SG, Tuske S, Tu X, Clark AD, Boyer PL, Hou X, Gaffney BL, Jones RA, Miller MD, Hughes SH, Arnold E. Structural Basis for the Role of the K65R Mutation in HIV-1 Reverse Transcriptase Polymerization, Excision Antagonism, and Tenofovir Resistance. Journal of Biological Chemistry. 2009;284(50):35092–100. doi: 10.1074/jbc.m109.022525.

52. Cilento ME, Ong YT, Tedbury PR, Sarafianos SG. Drug Interactions in Lenacapavir-Based Long-Acting Antiviral Combinations. Viruses. 2022;14(6):1202. doi: 10.3390/v14061202.

53. Michailidis E, Ryan EM, Hachiya A, Kirby KA, Marchand B, Leslie MD, Huber AD, Ong YT, Jackson JC, Singh K, Kodama EN, Mitsuya H, Parniak MA, Sarafianos SG. Hypersusceptibility mechanism of Tenofovir-resistant HIV to EFdA. Retrovirology. 2013;10(1):65. doi: 10.1186/1742-4690-10-65.

54. Snyder AA, Kaufman IL, Risener CJ, Kirby KA, Sarafianos SG. HIV-1 Reverse Transcriptase interactions with Long-acting NNRTI, Depulfavirine (VM1500A). 2026.

55. Huang H, Chopra R, Verdine GL, Harrison SC. Structure of a Covalently Trapped Catalytic Complex of HIV-1 Reverse Transcriptase: Implications for Drug Resistance. Science. 1998;282(5394):1669–75. doi: 10.1126/science.282.5394.1669.

56. Schneider DK, Soares AS, Lazo EO, Kreitler DF, Qian K, Fuchs MR, Bhogadi DK, Antonelli S, Myers SS, Martins BS, Skinner JM, Aishima J, Bernstein HJ, Langdon T, Lara J, Petkus R, Cowan M, Flaks L, Smith T, Shea-Mccarthy G, Idir M, Huang L, Chubar O, Sweet RM, Berman LE, McSweeney S, Jakoncic J. AMX – the Highly Automated Macromolecular Crystallography (17-ID-1) Beamline at the NSLS-II. Journal of Synchrotron Radiation. 2022;29(6):1480–94. doi: 10.1107/s1600577522009377.

57. Ralston CY, Gupta S, Del Mundo JT, Soe AC, Russell B, Rad B, Tyler J, Paul S, Kahan DN, Kristensen LG, Subramanian S, Kidd S, Burnett K, Sankaran B, Classen S, Prigozhin DM, Taylor JR, Dickert JM, Royal KB, Rozales A, Ortega SL, Allaire M, Nix JC, Hura GL, Holton JM, Hammel M, Adams PD. ALS-ENABLE: creating synergy and opportunity at the Advanced Light Source synchrotron structural biology beamlines. Journal of Synchrotron Radiation. 2025;32(4):1059–67. doi: 10.1107/s1600577525004205.

58. Kabsch W. *XDS*. Acta Crystallographica Section D Biological Crystallography. 2010;66(2):125-32. doi: 10.1107/s0907444909047337.

59. Zwart PH, Grosse-Kunstleve RW, Adams PD. Xtriage and Fest : automatic assessment of X-ray data and substructure structure factor estimation. 2005.

60. McCoy AJ, Grosse-Kunstleve RW, Adams PD, Winn MD, Storoni LC, Read RJ. *Phaser* rystallographic software. Journal of Applied Crystallography. 2007;40(4):658–74. doi: 10.1107/s0021889807021206.

61. Liebschner D, Afonine PV, Baker ML, Bunkóczi G, Chen VB, Croll TI, Hintze B, Hung L-W, Jain S, McCoy AJ, Moriarty NW, Oeffner RD, Poon BK, Prisant MG, Read RJ, Richardson JS, Richardson DC, Sammito MD, Sobolev OV, Stockwell DH, Terwilliger TC, Urzhumtsev AG, Videau LL, Williams CJ, Adams PD. Macromolecular structure determination using X-rays, neutrons and electrons: recent developments in *Phenix*. Acta Crystallographica Section D Structural Biology. 2019;75(10):861–77. doi: 10.1107/s2059798319011471.

62. Emsley P, Lohkamp B, Scott WG, Cowtan K. Features and development of *Coot*. Acta Crystallographica Section D Biological Crystallography. 2010;66(4):486-501. doi: 10.1107/s0907444910007493.

63. Chen VB, Arendall WB, Headd JJ, Keedy DA, Immormino RM, Kapral GJ, Murray LW, Richardson JS, Richardson DC. *MolProbity*: all-atom structure validation for macromolecular crystallography. Acta Crystallographica Section D Biological Crystallography. 2010;66(1):12-21. doi: 10.1107/s0907444909042073.

64. Morin A, Eisenbraun B, Key J, Sanschagrin PC, Timony MA, Ottaviano M, Sliz P. Collaboration gets the most out of software. eLife. 2013;2. doi: 10.7554/elife.01456.

65. Burley SK, Berman HM, Bhikadiya C, Bi C, Chen L, Costanzo LD, Christie C, Duarte JM, Dutta S, Feng Z, Ghosh S, Goodsell DS, Green RK, Guranovic V, Guzenko D, Hudson BP, Liang Y, Lowe R, Peisach E, Periskova I, Randle C, Rose A, Sekharan M, Shao C, Tao Y-P, Valasatava Y, Voigt M, Westbrook J, Young J, Zardecki C, Zhuravleva M, Kurisu G, Nakamura H, Kengaku Y, Cho H, Sato J, Kim JY, Ikegawa Y, Nakagawa A, Yamashita R, Kudou T, Bekker G-J, Suzuki H, Iwata T, Yokochi M, Kobayashi N, Fujiwara T, Velankar S, Kleywegt GJ, Anyango S, Armstrong DR, Berrisford JM, Conroy MJ, Dana JM, Deshpande M, Gane P, Gáborová R, Gupta D, Gutmanas A, Koča J, Mak L, Mir S, Mukhopadhyay A, Nadzirin N, Nair S, Patwardhan A, Paysan-Lafosse T, Pravda L, Salih O, Sehnal D, Varadi M, Vařeková R, Markley JL, Hoch JC, Romero PR, Baskaran K, Maziuk D, Ulrich EL, Wedell JR, Yao H, Livny M, Ioannidis YE. Protein Data Bank: the single global archive for 3D macromolecular structure data. Nucleic Acids Research. 2019;47(D1):D520–D8. doi: 10.1093/nar/gky949.

