## Supplemental Information for "Unraveling the Mechanism of HIV-1 Hypersusceptibility to Tenofovir Imparted by Islatravir Resistance Mutations"

### Supplemental Information (S.I.)

**Shreya M. Ravichandran[1,2], Alexa A. Snyder[1,2], Katelyn H. Kim[1,2], Isabella L. Kaufman[1,2], Xin Wen[1,2], Eleftherios Michailidis[1,2], Karen A. Kirby[1,2], Stefan G. Sarafianos*[1,2]**

[1]Center for ViroScience and Cure, Laboratory of Biochemical Pharmacology, Department of Pediatrics, Emory University School of Medicine, Atlanta, GA 30322

[2]Children's Healthcare of Atlanta, Atlanta, GA 30322

**KEYWORDS:** human immunodeficiency virus (HIV), acquired immunodeficiency syndrome (AIDS), islatravir, tenofovir, antiretroviral therapy (ART), reverse transcriptase, resistance, hypersusceptibility

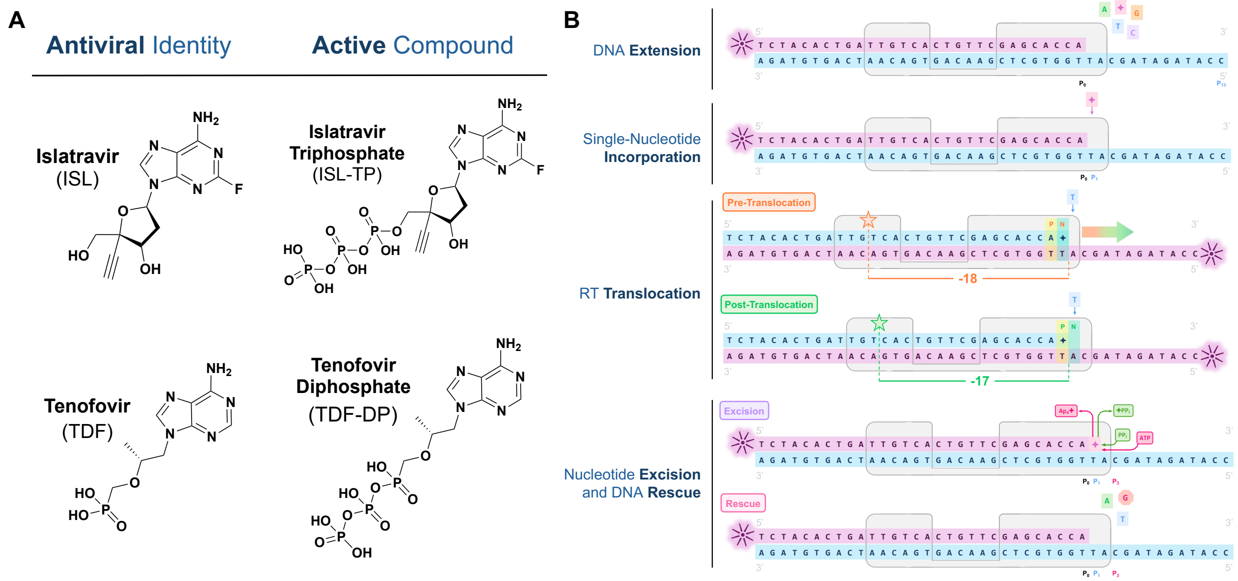


**Fig. S1. Structures of antivirals and steps of DNA synthesis studied in this work.** *[A]* Structures of ISL, TDF, and their phosphorylated forms. *[B]* During proviral DNA synthesis ("DNA Extension"), HIV-1 reverse transcriptase (RT) undergoes several steps that can each be targeted: incorporation of an incoming nucleotide onto the nascent strand ("Single-Nucleotide Incorporation"); translocation of the enzyme along the strand ("RT Translocation"); and – in response – resistance-driven excision of improper bases, such as inhibitory analogs, to “rescue” the elongating chain ("Nucleotide Excision and DNA Rescue"). ART-resistant or -hypersusceptible mutations can alter the inhibitors’ effects at any of these steps. Inhibitor: ✦; Ap4: adenosine tetraphosphate; dNTPs: A, T, G, C.

**DATA COLLECTION AND REFINEMENT STATISTICS**

| **Complex Identity** | **HIV-1 $RT_{M184V/A114S}$ / dsDNA / ISL-TP** | **HIV-1 $RT_{M184V}$ / dsDNA / ISL-TP** | **HIV-1 $RT_{A114S}$ / dsDNA / ISL-TP** |
|---|---|---|---|
| **PDB ID** | **12SK** | **12SM** | **12SR** |
| **Data Collection and Processing** | | | |
| Structural Method | X-Ray Crystallography | X-Ray Crystallography | X-Ray Crystallography |
| Synchrotron Beamline | NSLS-II AMX (17-ID-1) | ALS 4.2.2 | NSLS-II AMX (17-ID-1) |
| Wavelength (nm) | 0.979338 | 1.000075 | 0.919491 |
| Processing Program | XDS | XDS | XDS |
| Resolution Range (Å) | 33.98 – 2.79 | 53.54 – 2.91 | 33.68 – 2.65 |
| Space Group | C 2 2 $2_1$ | C 2 2 $2_1$ | C 2 2 $2_1$ |
| Unit Cell Constants *a, b, c* | 168.97Å 171.08Å 106.27Å | 167.59Å 169.50Å 101.68Å | 167.52Å 170.56Å 105.31Å |
| Total Reflections | 504,864 (80,806) | 197,860 (21,642) | 421,910 (66,706) |
| Unique Reflections | 37,252 (5,941) | 56,497 (7,549) | 43,906 (6,752) |
| Multiplicity | 13.6 (13.6) | 3.5 (2.9) | 9.6 (9.9) |
| Data Completeness (%) | 96.7 (97.2) | 91.8 (76.3) | 99.1 (96.1) |
| Mean I/σ(I) | 15.8 (2.4) | 11.3 (1.5) | 14.5 (1.8) |
| Wilson B-factor ($Å^2$) | 62.2 | 60.4 | 68.7 |
| $R_{meas}$ | 0.16 (1.2) | 0.11 (1.1) | 0.12 (1.1) |
| $CC_{1/2}$ | 0.998 (0.801) | 0.996 (0.651) | 0.998 (0.782) |
| **Data Refinement** | | | |
| Refinement Program | PHENIX 2.0_5885 | PHENIX 2.0_5885 | PHENIX 2.0_5885 |
| No. of Reflections (Working) | 37,224 | 30,215 | 43,875 |
| No. of Reflections (Test) | 1,989 (5.3%) | 1,538 (5.1%) | 2,267 (5.2%) |
| $R_{work}$ | 0.195 | 0.203 | 0.227 |
| $R_{free}$ | 0.251 | 0.259 | 0.255 |
| Total Number of Atoms | 8,917 | 8,847 | 8,944 |
| Average B, all atoms ($Å^2$) | 57.7 | 62.1 | 70.4 |
| Ligand PDB IDs | 6FN, MRG | 6FN, MRG | 6FN, MRG |
| **Validation Statistics** | | | |
| MolProbity Score | 1.18 | 1.37 | 1.04 |
| Clashscore | 3.81 | 5.01 | 2.51 |
| Ramachandran Plot | | | |
| Favored (%) | 98.02 | 98.54 | 98.31 |
| Allowed (%) | 1.77 | 1.36 | 1.69 |
| Disallowed (%) | 0.21 | 0.10 | 0.00 |
| Bond Lengths RMSD (Å) | 0.003 | 0.002 | 0.002 |
| Bond Angles RMSD (°) | 0.526 | 0.497 | 0.471 |

**Table S1. Data collection and refinement statistics for ISL-TP-containing structures.** Values reported for PDB IDs 12SK ($RT_{M184V/A114S}$), 12SR ($RT_{M184V}$), and 12SM ($RT_{A114S}$).

**DATA COLLECTION AND REFINEMENT STATISTICS**

| **Complex Identity** | **HIV-1 $RT_{M184V/A114S}$ / dsDNA / TDF-DP** |
|---|---|
| **PDB ID** | **12SW** |
| **Data Collection and Processing** | |
| Structural Method | X-Ray Crystallography |
| Synchrotron Beamline | NSLS-II AMX (17-ID-1) |
| Wavelength (nm) | 0.919491 |
| Processing Program | XDS |
| Resolution Range (Å) | 33.73 – 3.20 |
| Space Group | C 2 2 $2_1$ |
| Unit Cell Constants *a, b, c* | 165.71Å 172.25Å 105.42Å |
| Total Reflections | 114,137 (18,180) |
| Unique Reflections | 25,051 (3,921) |
| Multiplicity | 4.6 (4.6) |
| Data Completeness (%) | 99.4 (98.2) |
| Mean I/σ(I) | 7.3 (1.6) |
| Wilson B-factor ($Å^2$) | 91.6 |
| $R_{meas}$ | 0.18 (1.2) |
| $CC_{1/2}$ | 0.995 (0.490) |
| **Data Refinement** | |
| Refinement Program | PHENIX 2.0_5885 |
| No. of Reflections (Working) | 25,051 |
| No. of Reflections (Test) | 1,226 (4.9%) |
| $R_{work}$ | 0.208 |
| $R_{free}$ | 0.243 |
| Total Number of Atoms | 8,724 |
| Average B, all atoms ($Å^2$) | 82.0 |
| Ligand PDB IDs | TNV, MRG |
| **Validation Statistics** | |
| MolProbity Score | 1.60 |
| Clashscore | 5.68 |
| Ramachandran Plot | |
| Favored (%) | 97.15 |
| Allowed (%) | 2.74 |
| Disallowed (%) | 0.11 |
| Bond Lengths RMSD (Å) | 0.003 |
| Bond Angles RMSD (°) | 0.582 |

| **Complex Identity** | **HIV-1 $RT_{M184V}$ / dsDNA / TDF-DP** | **HIV-1 $RT_{A114S}$ / dsDNA / TDF-DP** |
|---|---|---|
| **PDB ID** | **12TG** | **12TK** |
| **Data Collection and Processing** | | |
| Structural Method | Single-Particle Cryo-EM | Single-Particle Cryo-EM |
| Collection Facility | BNL LBMS | Emory University IEMC |
| Microscope | TFS Krios Cryo-TEM | TFS Talos Arctica Cryo-TEM |
| Electron Detector | Gatan K3 | Gatan K3 BioQuantum |
| Magnification (X) | 105,000 | 100,000 |
| Voltage (kV) | 300 | 200 |
| Electron Exposure ($e^-/Å^2$) | 50.0 | 50.5 |
| Exposure Time per Image (s) | 2.27 | 3.04 |
| Defocus Range (μm) | -2.0 to -1.0 | -2.2 to -0.8 |
| Pixel Size (Å/pix) | 0.4125 | 0.84 |
| Processing Program | CryoSPARC 4.7.1 | CryoSPARC 4.7.1 |
| Initial No. of Micrographs | 4,637 | 3,108 |
| Final No. of Micrographs | 3,931 | 2,815 |
| Initial No. of Particles | 2,663,102 | 4,049,858 |
| Final No. of Particles | 665,680 | 572,304 |
| Global Map Resolution (Å) | 2.63 | 2.87 |
| FSC Threshold | 0.143 | 0.143 |
| Map Contour Level | 0.01 | 0.025 |
| **Data Refinement** | | |
| Refinement Program | PHENIX 2.0_5885 | PHENIX 2.0_5885 |
| Total Number of Atoms | 8,754 | 8,822 |
| Ligand PDB IDs | TNV, MRG | TNV, MRG |
| EMDB IDs | EMD-76741 | EMD-76742 |
| **Validation Statistics** | | |
| MolProbity Score | 1.49 | 1.02 |
| Clashscore | 5.02 | 1.33 |
| $CC_{mask}$ | 0.85 | 0.88 |
| Ramachandran Plot | | |
| Favored (%) | 98.21 | 97.61 |
| Allowed (%) | 1.68 | 2.39 |
| Disallowed (%) | 0.11 | 0.00 |
| Bond Lengths RMSD (Å) | 0.002 | 0.002 |
| Bond Angles RMSD (°) | 0.453 | 0.428 |

**Table S2. Data collection and refinement statistics for TDF-DP-containing structures.** Values reported for PDB IDs 12SW ($RT_{M184V/A114S}$), 12TG ($RT_{M184V}$), and 12TK ($RT_{A114S}$) / EMDB IDs EMD-76741 ($RT_{M184V}$) and EMD-76742 ($RT_{A114S}$).

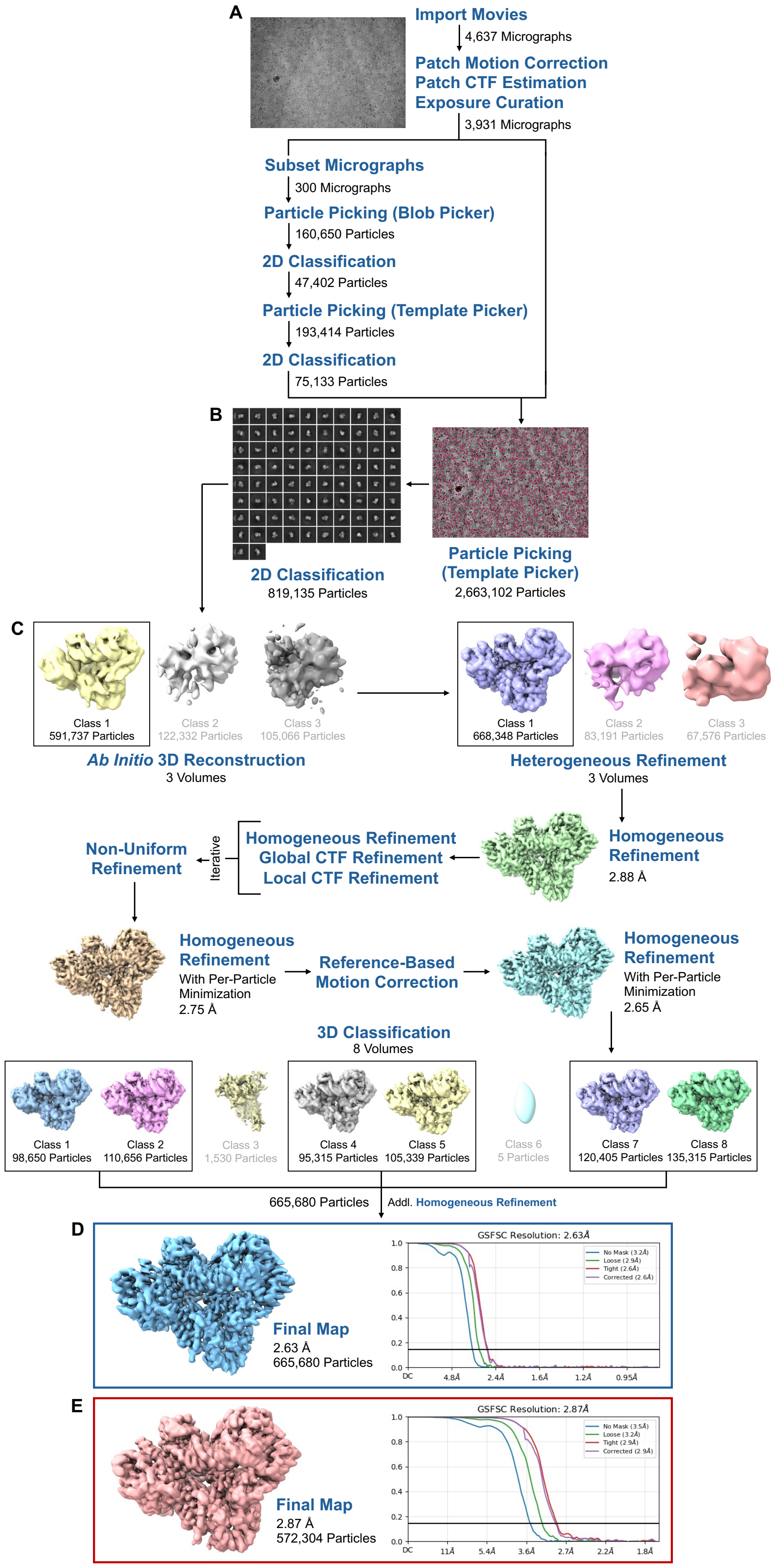


**Fig. S2. Dataset processing workflows for cryo-EM structures.** *[A]* To obtain high-quality particles of the $RT_{M184V}$/dsDNA/TDF-DP ternary complex, a subset of 300 micrographs (out of 4,637 total) was used for initial particle extraction and 2D classification jobs. *[B]* 75,133 subset-derived particles were then used as 2D templates to extract and classify particles from 3,931 curated movies. *[C]* From an *ab initio* volume, multiple refinement jobs (homogeneous, heterogeneous, global CTF, local CTF, non-uniform) – along with Reference-Based Motion Correction and 3D Classification – were carried out to improve the map. *[D]* The resulting map of $RT_{M184V}$/dsDNA/TDF-DP (blue) was determined to a global resolution of 2.63 Å. *[E]* A highly similar workflow was conducted for the $RT_{A114S}$/dsDNA/TDF-DP ternary complex, resulting in a map (salmon) determined to 2.87 Å global resolution (2,815/3,107 micrographs selected; 572,304 particles used). All jobs conducted in CryoSPARC v4.7.1 (Structura Biotechnology, Inc.).

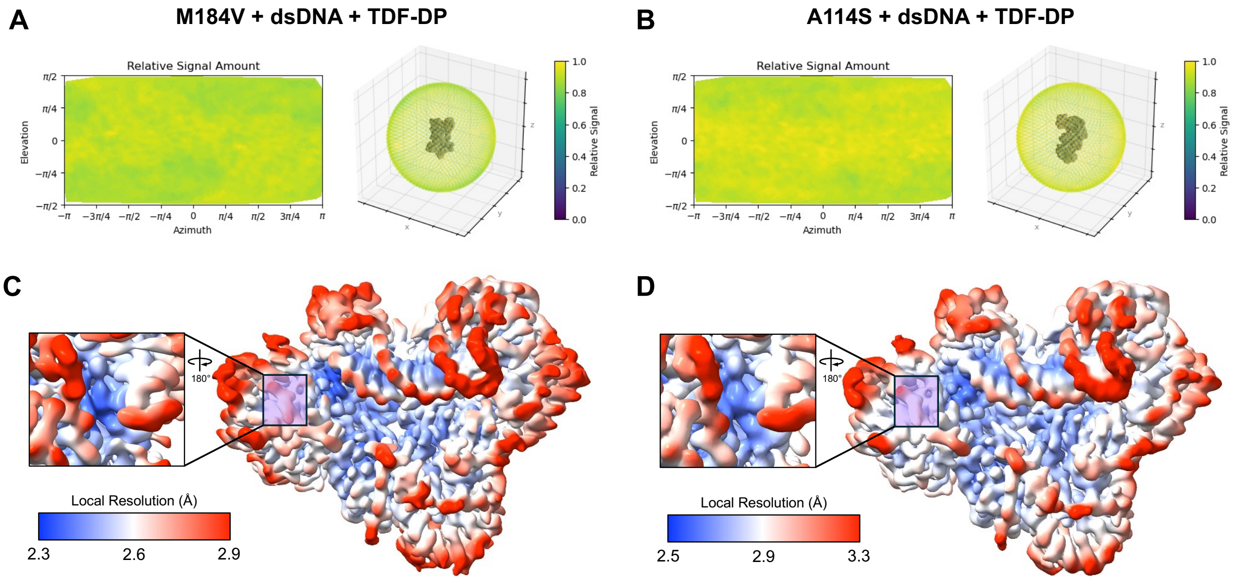


**Fig. S3. Sampling of particle orientations and local resolution maps of cryo-EM structures.** *[A-B]* Orientation Diagnostics jobs were used to show distributions of captured particle orientations for both *[A]* $RT_{M184V}$ and *[B]* $RT_{A114S}$ ternary complexes. *[C-D]* Local Resolution Estimation jobs were used to identify local resolution ranges at the polymerase active site (inset, purple box) in both cryo-EM structures. *[C]* $RT_{M184V}$/dsDNA/TDF-DP resolution range: 2.3 Å (blue) – 2.9 Å (red); *[D]* $RT_{A114S}$/dsDNA/TDF-DP resolution range: 2.5 Å (blue) – 3.3 Å (red). All jobs conducted in CryoSPARC v4.7.1 (Structura Biotechnology, Inc.). Local resolution maps displayed using UCSF ChimeraX v1.11.

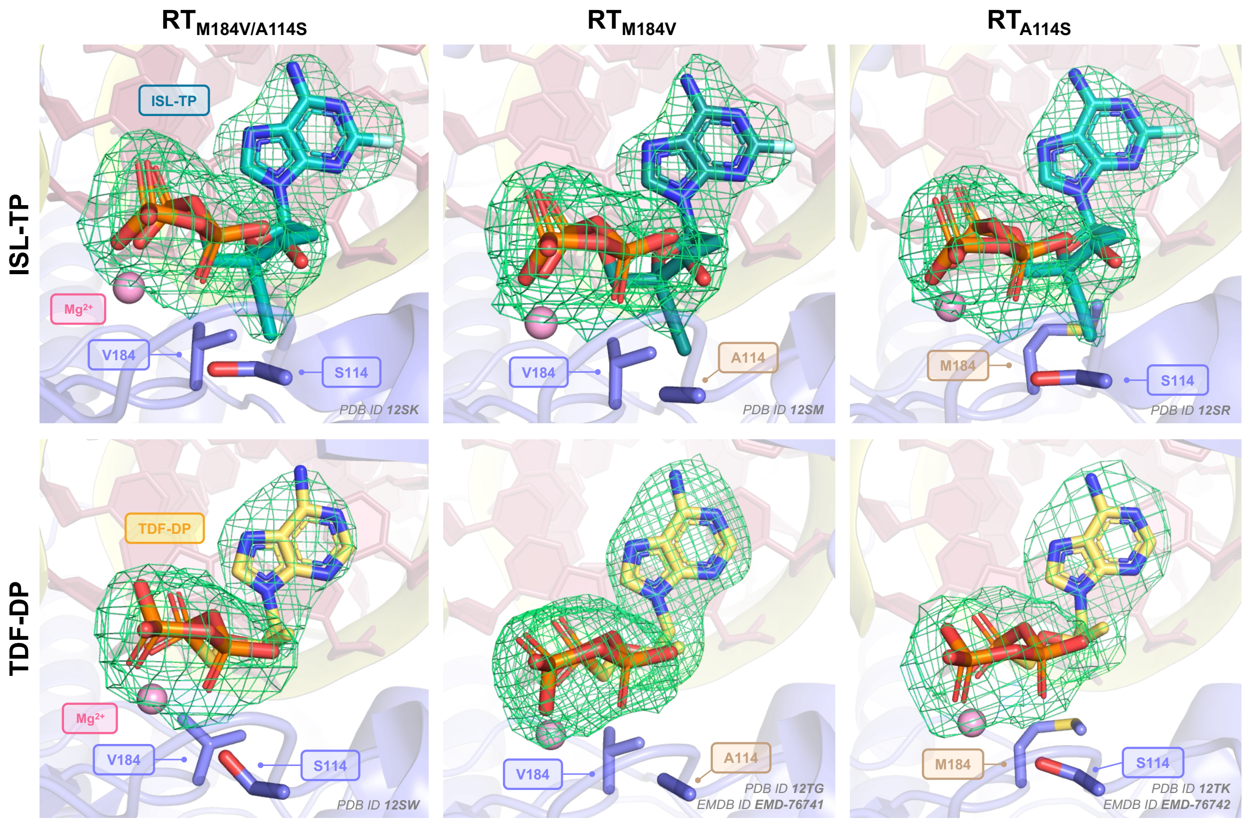


**Fig. S4. Ligand density maps across complexes.** For all crystal structures, $2F_o$-$F_c$ difference map is displayed at σ = 1.000 RMSD / map level = 0.167 eV/A$^3$. For each cryo-EM structure, Coulombic density map is displayed at the respective map contour level ($RT_{M184V}$ + TDF-DP: 0.01; $RT_{A114S}$ + TDF-DP: 0.025). Color palette: RT: purple; dsDNA backbone: yellow; dsDNA bases: pink; ISL-TP: teal; TDF-DP: yellow; $Mg^{2+}$ ion: light pink; phosphorus: orange; oxygen: red; nitrogen: blue; sulfur: yellow; density mesh: green; WT residue labels: brown; mutant residue labels: purple. PDB ID (and EMDB ID, as applicable) listed in bottom right corner of each panel. Density maps generated in UCSF ChimeraX v1.11 using ISOLDE v1.10 (Altos Labs, Inc.). Figure created in PyMOL v3.1.3 (Schrödinger, LLC.).

| | |
|---|---|
| **Primer Extension, Incorporation, and Excision** | |
| $P_{d18}$-$P_0$ | 5’ – † GTC ACT GTT CGA GCA CCA – 3’ |
| $T_{d31}$ | 3’ – CAG TGA CAA GCT CGT GGT TAC GAT CGA TAC C – 5' |
| **Hydroxyl-Radical DNA Footprinting** | |
| $P_{d30}$-$P_0$ | 5’ – TCT ACA CTG ATT GTC ACT GTT CGA GCA CCA – 3’ |
| $T_{d43}$ | 3’ – AGA TGT GAC TAA CAG TGA CAA GCT CGT GGT TAC GAT AGA TAC C † – 5' |
| **X-Ray Crystallography and Cryo-EM** | |
| $P_{d18}$* | 5’ – GTC CCT GTT CGG ‡GCG CCG – 3’ |
| $T_{d24}$ | 3’ – GT CAG GGA CAA GCC CGC GGC TGG T – 5’ |

**Table S3. Oligonucleotide sequences used for *in vitro* assays.** For all mechanistic studies, the dsDNA substrate possessed one Cyanine 3 (Cy3) fluorophore-labeled DNA strand to be tracked for subsequent gel quantification. Primer sequences for biochemical studies were designed based on the previously characterized $P_0$ site. For structural studies, a previously characterized dsDNA substrate was used for RT-DNA crosslinking through addition of a thioalkyl group on a guanosine within the primer strand. In all assays, a 3:1 molar ratio of labeled to non-labeled strand was used to ensure complete annealing of the labeled strand. †Indicates location of Cy3 fluorophore. ‡Indicates location of thioalkyl modification.

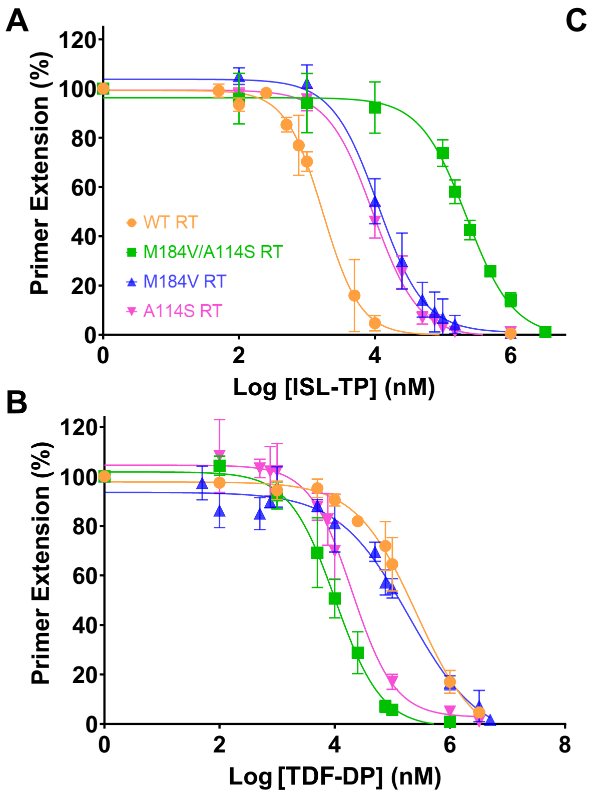


C

| HIV-1 RT Variant | Antiviral | *Half-Max. Extension Concentration* (μM) [a] |
|---|---|---|
| WT | ISL-TP | 1.86 ± 0.67 |
| | TDF-DP | 270 ± 129 |
| M184V/ A114S | ISL-TP | 216 ± 33.9 |
| | TDF-DP | 10.5 ± 3.95 |
| M184V | ISL-TP | 11.9 ± 3.38 |
| | TDF-DP | 188 ± 84.4 |
| A114S | ISL-TP | 9.65 ± 1.72 |
| | TDF-DP | 18.8 ± 7.45 |

**Fig. S5. *In vitro* ISL-TP and TDF-DP inhibition profiles during HIV-1 RT primer extension.** *[A]* Results of ISL-TP-based inhibition of Cy3-labeled DNA polymerization among the RT variants. *[B]* Results of TDF-DP-based inhibition of Cy3-labeled DNA polymerization among the RT variants. *[C]* Inhibitor concentrations leading to half-maximal inhibition of primer extension are reported. Color palette: $RT_{WT}$: orange; $RT_{M184V/A114S}$: green; $RT_{M184V}$: blue; $RT_{A114S}$: pink. Averages and errors calculated from three experiments. *[a]* Concentrations were reported with respective standard errors based on four-parameter inhibition, with calculations and graphs done using GraphPad Prism 9.

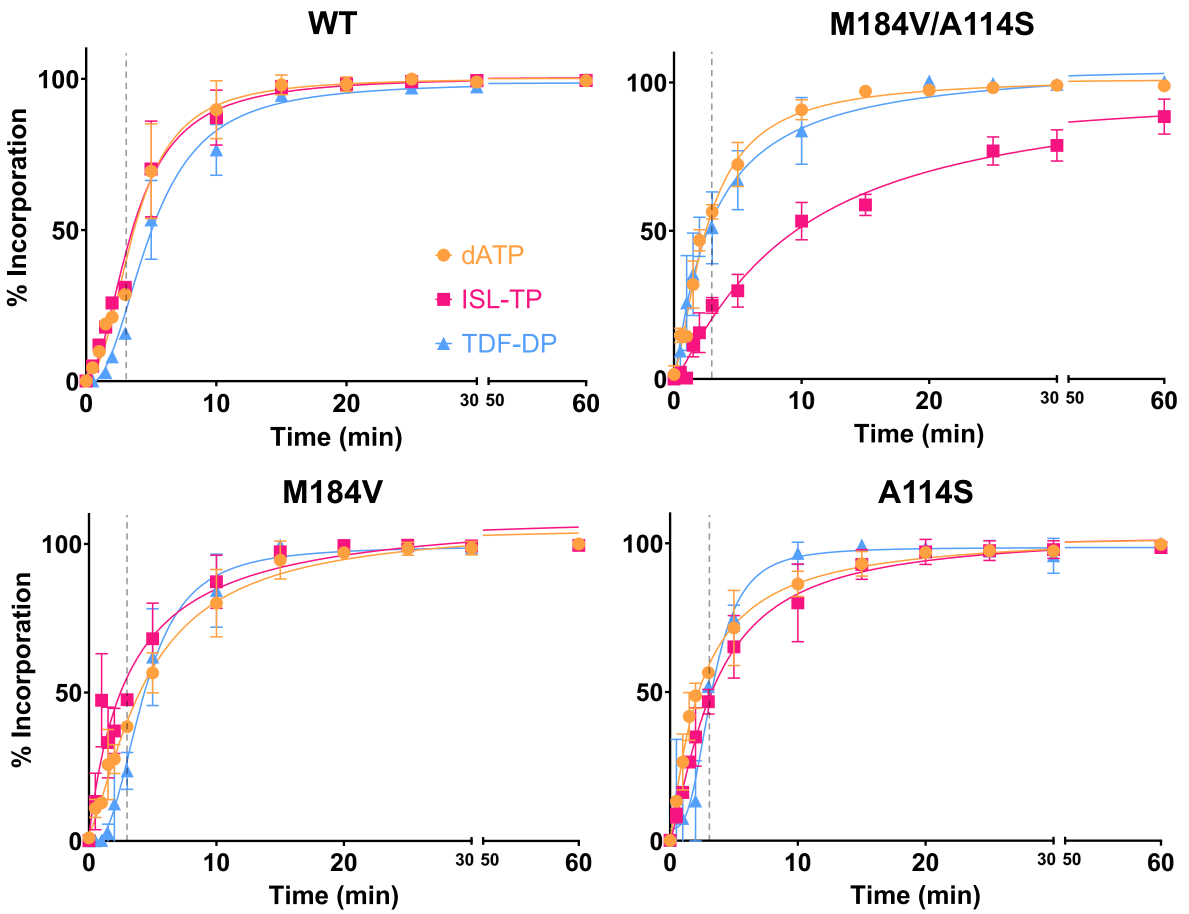


**Fig. S6. Time-courses of nucleotide and analog incorporation among HIV-1 RT mutants.** In determining a timepoint across all RT-nucleotide conditions at which a linear incorporation rate occurs, time-dependent incorporation assays were conducted up to 1 h. Three-min timepoint highlighted as dashed gray line. An x-axis break was added between 30 and 50 m. Color palette: dATP: light orange; ISL-TP: light pink; TDF-DP: light blue. Error bars calculated from two-to-four experiments. Graphs created in GraphPad Prism 9.

| RT Mutant | dNTP | $K_m$ (μM) [A] | $k_{cat}$ ($min^{-1}$) | $k_{cat}/K_m$ ($min^{-1}$ $μM^{-1}$) |
|---|---|---|---|---|
| **Wild-Type** | **dATP** | 0.25 ± 0.05 | 2.44 ± 0.10 | 9.76 ± 1.85 |
| | **ISL-TP** | 0.18 ± 0.04 | 2.22 ± 0.09 | 12.3 ± 2.64 |
| | **TDF-DP** | 0.94 ± 0.14 | 1.81 ± 0.07 | 1.93 ± 0.29 |
| **M184V/ A114S** | **dATP** | 0.31 ± 0.06 | 2.27 ± 0.10 | 7.32 ± 1.53 |
| | **ISL-TP** | 43 ± 24 | 1.80 ± 0.67 | 0.04 ± 0.03 |
| | **TDF-DP** | 1.44 ± 0.21 | 2.74 ± 0.11 | 1.90 ± 0.28 |
| **M184V** | **dATP** | 0.11 ± 0.02 | 1.67 ± 0.09 | 15.2 ± 3.38 |
| | **ISL-TP** | 0.21 ± 0.04 | 2.09 ± 0.06 | 9.95 ± 1.83 |
| | **TDF-DP** | 1.22 ± 0.26 | 1.87 ± 0.10 | 1.53 ± 0.34 |
| **A114S** | **dATP** | 0.27 ± 0.05 | 1.99 ± 0.07 | 7.37 ± 1.31 |
| | **ISL-TP** | 0.35 ± 0.10 | 1.55 ± 0.09 | 4.43 ± 1.28 |
| | **TDF-DP** | 0.36 ± 0.06 | 1.56 ± 0.06 | 4.33 ± 0.79 |

**Table S4. Michaelis-Menten kinetics values from steady-state inhibitor incorporation assays.** *[A]* Values reported from three-to-five experiments. All values are reported with respective standard errors. Non-linear regression calculations performed under "Michaelis-Menten Kinetics" in GraphPad Prism 9. For catalytic efficiencies, propagated standard errors were calculated as described in Materials and Methods.

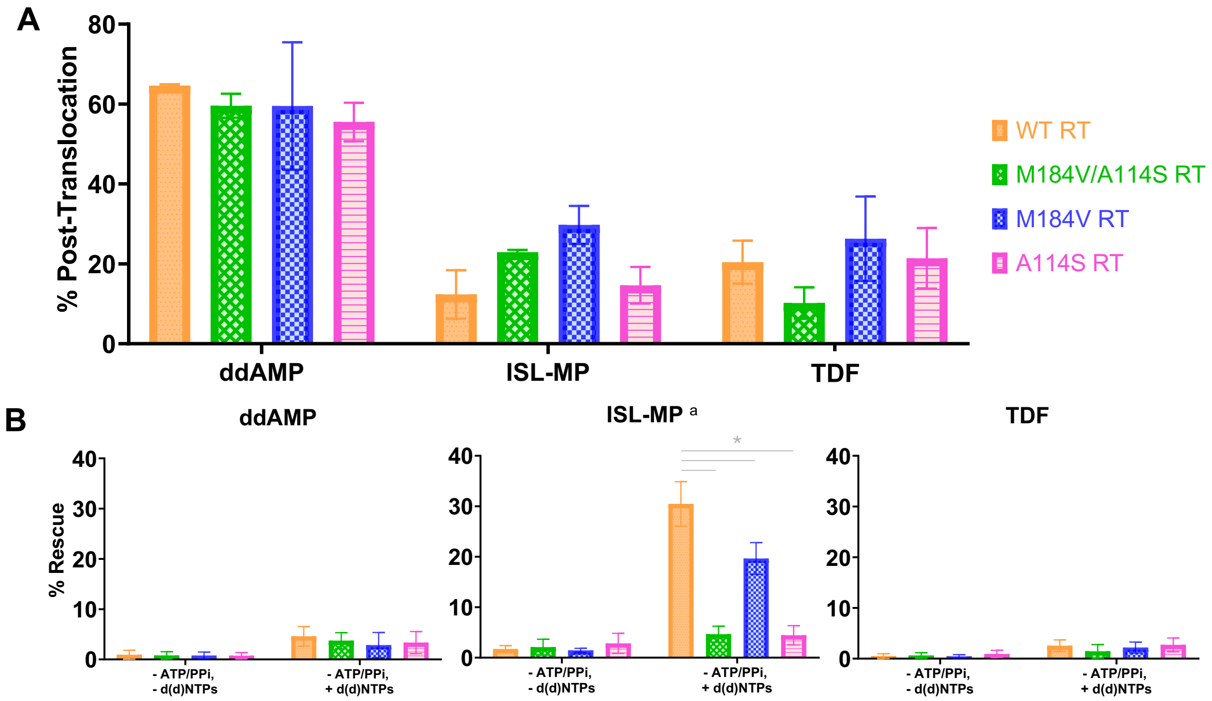


**Fig. S7. Negative controls for HIV-1 RT translocation and inhibitor excision / DNA rescue among RT mutants.** *[A]* The observed frequencies of post-translocated RT complexes are reported in the absence of the subsequent nucleotide, dTTP. *[B]* DNA rescue was assessed among the RT-antiviral pairings in the absence of excision-mediating ATP or $PP_i$. *[a]* Significant "discoveries" were identified from t-tests with Benjamini-Hochberg (BH) corrections to control for False Discovery Rate (FDR). Only comparisons marked as "discoveries" at $Q = 0.01$ were denoted with an asterisk (*). ISL-MP discoveries from BH-controlled t-tests: $RT_{WT}$ *vs.* $RT_{M184V/A114S}$ (mean diff = 25.80%; $q = 10^{-6}$; $p = 10^{-6}$; $df = 8$); $RT_{WT}$ *vs.* $RT_{M184V}$ (mean diff = 10.81%; $q = 1.31 \times 10^{-3}$; $p = 1.297 \times 10^{-3}$; $df = 8$); $RT_{WT}$ *vs.* $RT_{A114S}$ (mean diff = 26.02%; $q = 10^{-6}$; $p = 10^{-6}$; $df = 8$). Color palette: $RT_{WT}$: orange; $RT_{M184V/A114S}$: green; $RT_{M184V}$: blue; $RT_{A114S}$: pink. Averages and errors calculated from two-to-six experiments. Graphs created in GraphPad Prism 9.

| RT Mutant | dNTP | Translocation Controls<br>(-) dTTP | Nucleotide Excision and DNA Rescue Controls<br>(-) ATP / $PP_i$<br>(-) d(d)NTPs | <br>(-) ATP / $PP_i$<br>(+) d(d)NTPs |
|---|---|---|---|---|
| **Wild-Type** | **ddAMP** | 64.6 ± 0.33 | 0.92 ± 0.89 | 4.61 ± 1.95 |
| | **ISL-MP** | 12.3 ± 6.08 | 1.72 ± 0.68 | 30.5 ± 4.41 |
| | **TDF** | 20.4 ± 5.39 | 0.50 ± 0.46 | 2.56 ± 1.15 |
| **M184V/ A114S** | **ddAMP** | 59.6 ± 3.00 | 0.80 ± 0.72 | 3.73 ± 1.60 |
| | **ISL-MP** | 22.9 ± 0.59 | 2.09 ± 1.56 | 4.65 ± 1.59 |
| | **TDF** | 10.2 ± 3.93 | 0.63 ± 0.55 | 1.44 ± 1.35 |
| **M184V** | **ddAMP** | 59.5 ± 15.9 | 0.77 ± 0.70 | 2.86 ± 2.51 |
| | **ISL-MP** | 29.8 ± 4.75 | 1.46 ± 0.40 | 19.6 ± 3.15 |
| | **TDF** | 26.3 ± 10.5 | 0.46 ± 0.35 | 2.20 ± 1.08 |
| **A114S** | **ddAMP** | 55.5 ± 4.82 | 0.73 ± 0.63 | 3.35 ± 2.20 |
| | **ISL-MP** | 14.6 ± 4.57 | 2.84 ± 1.97 | 4.44 ± 1.90 |
| | **TDF** | 21.4 ± 7.58 | 0.93 ± 0.71 | 2.72 ± 1.35 |

**Table S5. Negative controls for HIV-1 RT translocation and inhibitor excision / DNA rescue among RT mutants.** Averages and standard deviations are reported for the controls of translocation and excision tests. Values reported from two-to-six experiments.

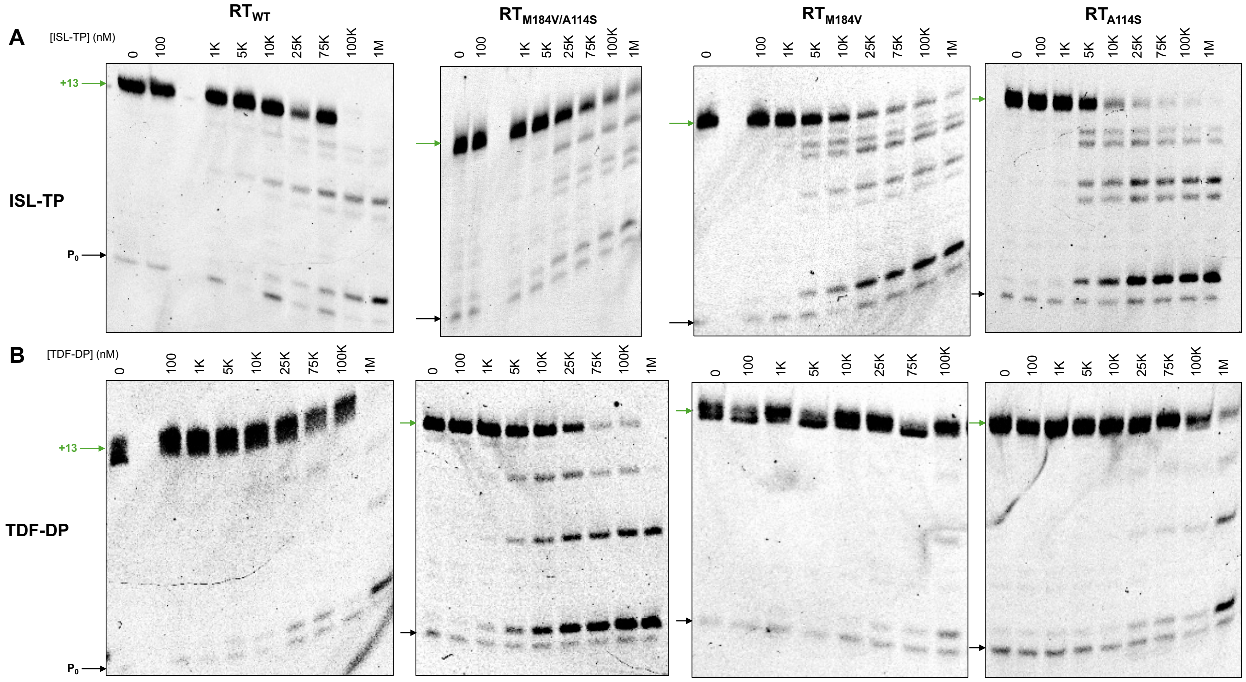


**Fig. S8. Representative gel images for results of *in vitro* inhibitor susceptibility during DNA primer extension.** Successful primer extension at each inhibitor concentration was calculated as the percentage of total signal in each lane obtained from the fully extended primer band (+13, green) beginning at the P$_0$ site (black). P$_{d18}$-Cy3 fluorescent signal was quantified using AzureSpot Pro v1.2.

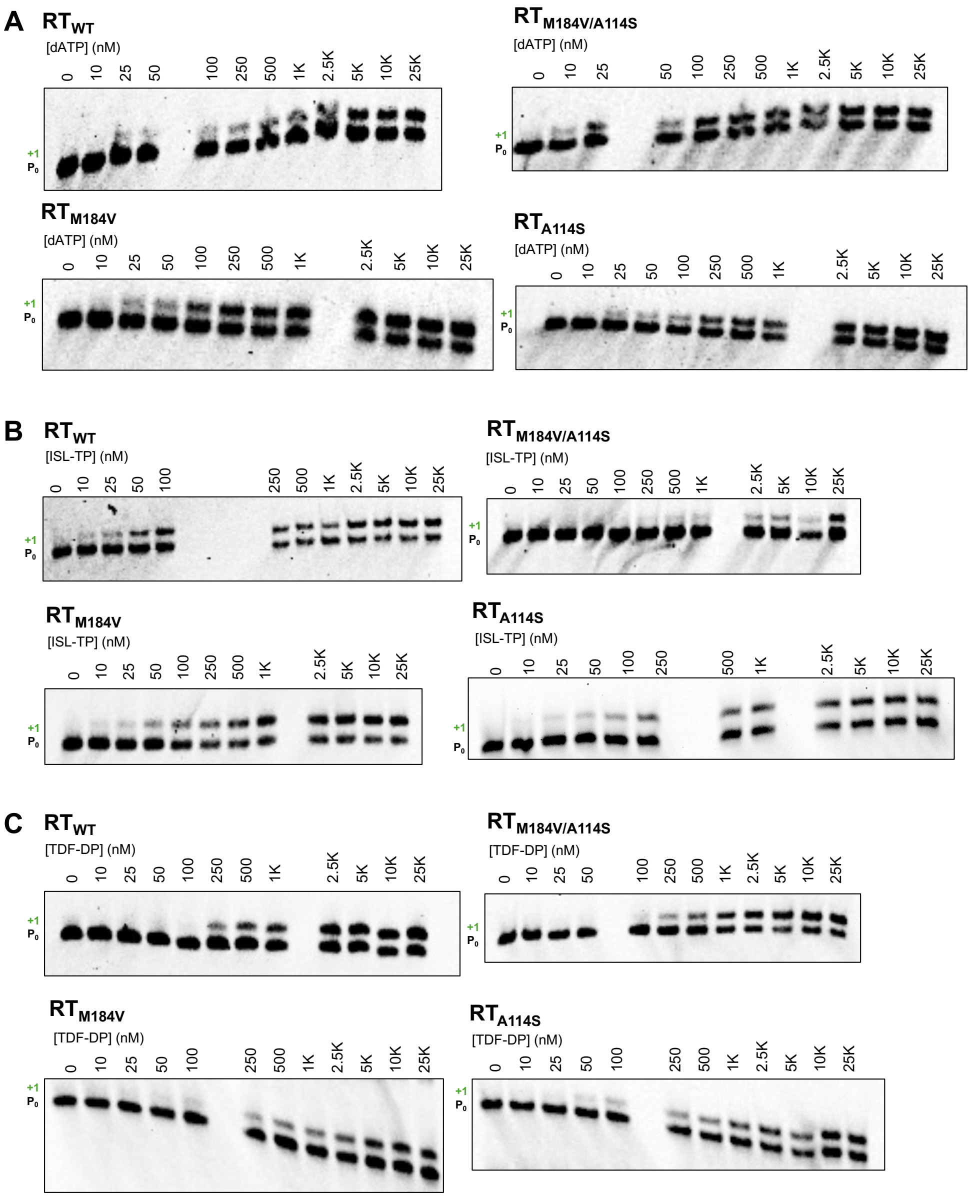


**Fig. S9. Representative gel images for results of concentration-dependent single-nucleotide incorporation.** *[A]* Concentration-dependent incorporation is shown for *[A]* dATP, *[B]* ISL-TP, and *[C]* TDF-DP. $P_{d18}$-Cy3 fluorescent signal was quantified using AzureSpot Pro v1.2.

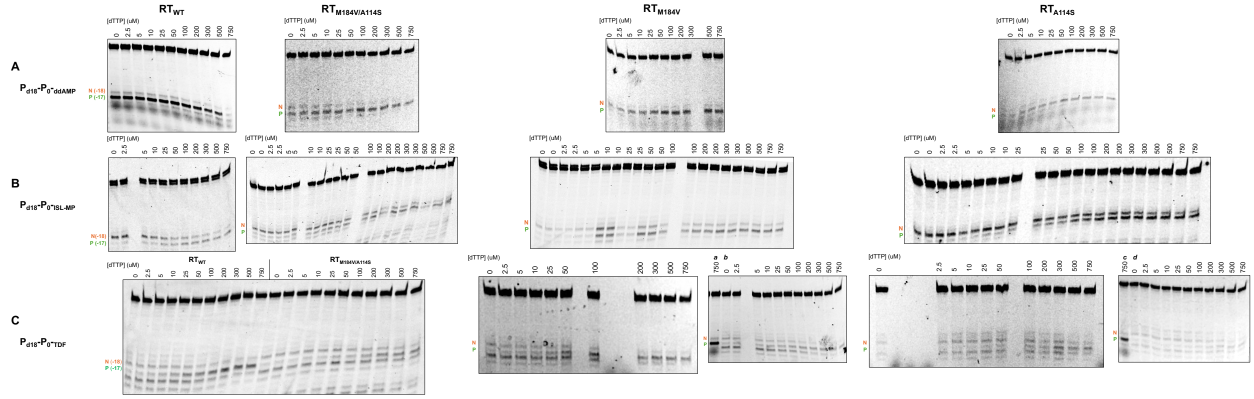

**Fig. S10. Representative gel images for results of hydroxyl radical iron footprinting.** Positioning of RT along the DNA substrate is measured as the ratio of enzyme observed at the *N*-site (orange; hydroxyl radical cleavage site located 18 bases away from position of inhibitor) to *P*-site (green; cleavage site located 17 bases away from position of inhibitor). For TDF gels, the *N*-site and *P*-site bands were identified in control gels: *[a]* $RT_{WT}$ + $T_{d43}$/$P_{d18}$-$P_0$-ddAMP + 750 µM dTTP; *[b]* $RT_{M184V}$ + $T_{d43}$/$P_{d18}$-$P_0$-TDF + dTTP; *[c]* $RT_{M184V/A114S}$ + $T_{d43}$/$P_{d18}$-$P_0$-ddAMP + 750 µM dTTP; *[d]* $RT_{A114S}$ + $T_{d43}$/$P_{d18}$-$P_0$-TDF + dTTP. $T_{d43}$-Cy3 fluorescent signal was quantified using AzureSpot Pro v1.2.

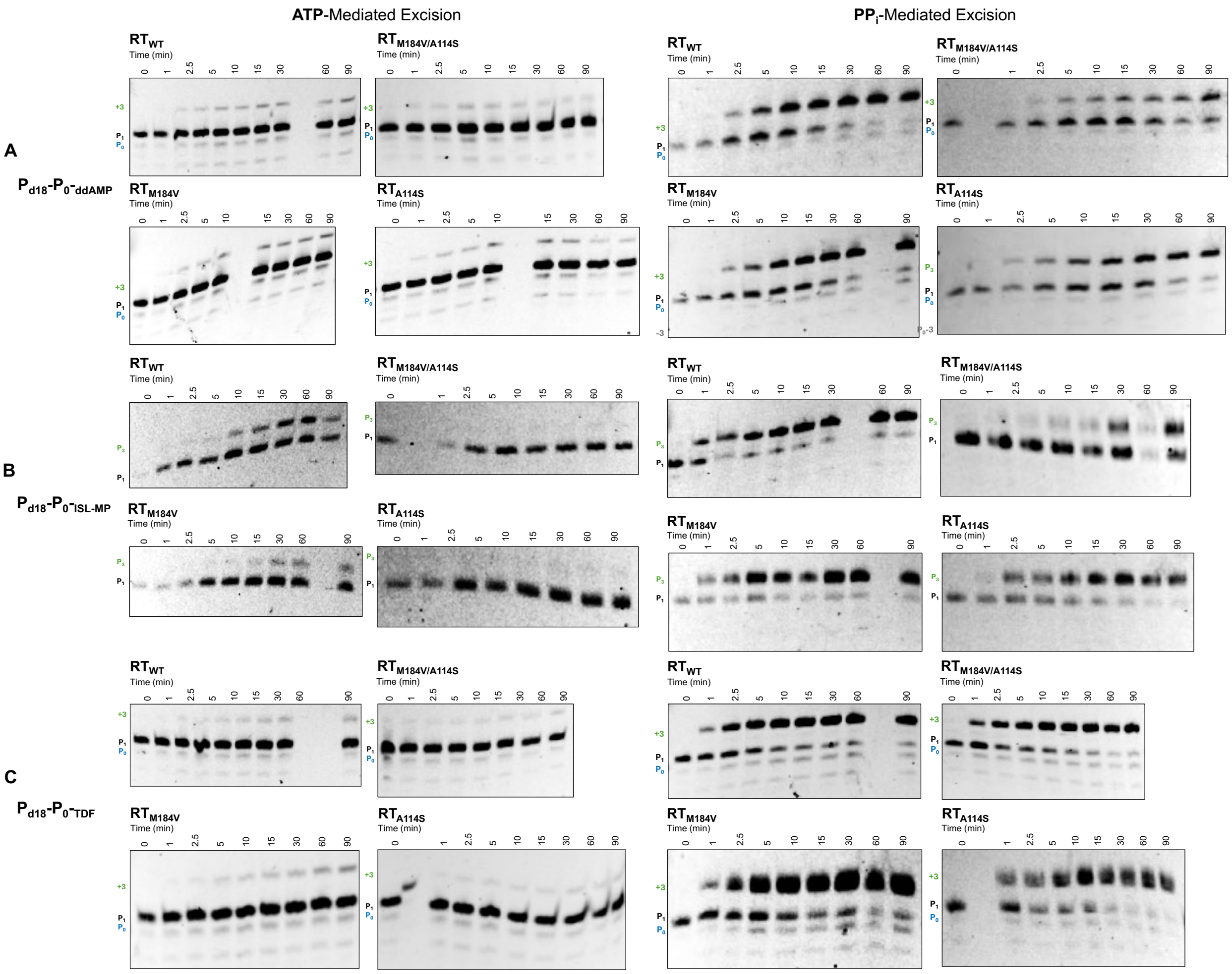


**Fig. S11. Representative gel images for results of ATP- and $PP_i$-mediated inhibitor excision.** Results shown for *[A]* ddAMP-terminated primer; *[B]* ISL-MP-terminated primer; *[C]* TDF-terminated primer. For quantification, the ratio of $P_1$ to $P_3$ was solely measured to assess the paired efficiency of DNA excision and rescue. $P_{d18}$-Cy3 fluorescent signal was quantified using AzureSpot Pro v1.2.

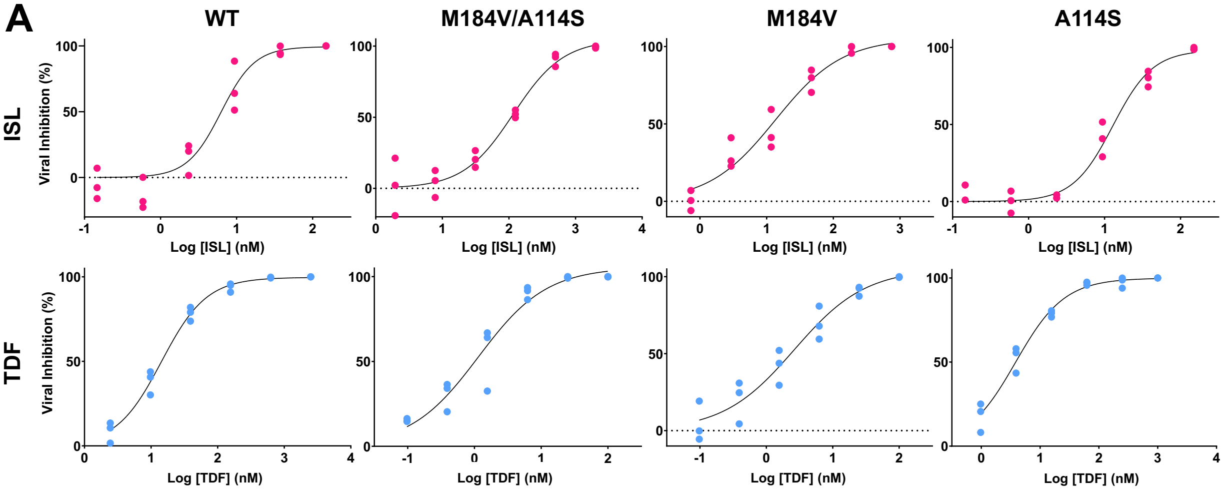


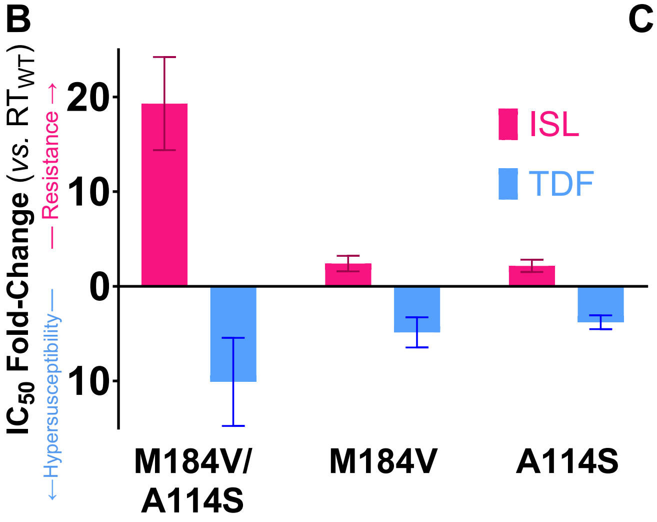


C

| VSV-G Pseudotyped Virus | Antiviral | $[IC_{50}]$ (nM) [a] | $IC_{50}$ Fold-Change (*vs.* WT) [b] |
|---|---|---|---|
| WT | ISL | 6.05 ± 2.54 | - |
| | TDF | 14.4 ± 3.48 | - |
| M184V/ A114S | ISL | 117 ± 15.8 | ↑ 19.3 ± 4.92 |
| | TDF | 1.43 ± 1.09 | ↓ 10.1 ± 4.66 |
| M184V | ISL | 14.5 ± 6.07 | ↑ 2.40 ± 0.82 |
| | TDF | 2.97 ± 1.53 | ↓ 4.87 ± 1.60 |
| A114S | ISL | 13.0 ± 4.11 | ↑ 2.16 ± 0.65 |
| | TDF | 3.80 ± 0.86 | ↓ 3.80 ± 0.73 |

**Fig. S12. *In cellulo* ISL and TDF profiles reported for pseudotyped HIV-1 variants.** *[A]* Inhibition curves against VSV-G pseudotyped viral mutants show respective $IC_{50}$s of virus-antiviral combinations (ISL: pink; TDF: blue). *[B-C]* Fold-changes in $IC_{50}$ values between mutant and WT pseudotyped viruses are reported. *[a-b]* $IC_{50}$s and fold-changes are reported with standard errors as described in Materials and Methods. Concentrations were reported with respective standard deviations from three experiments based on four-parameter inhibition, with calculations and graphs done using GraphPad Prism 9.

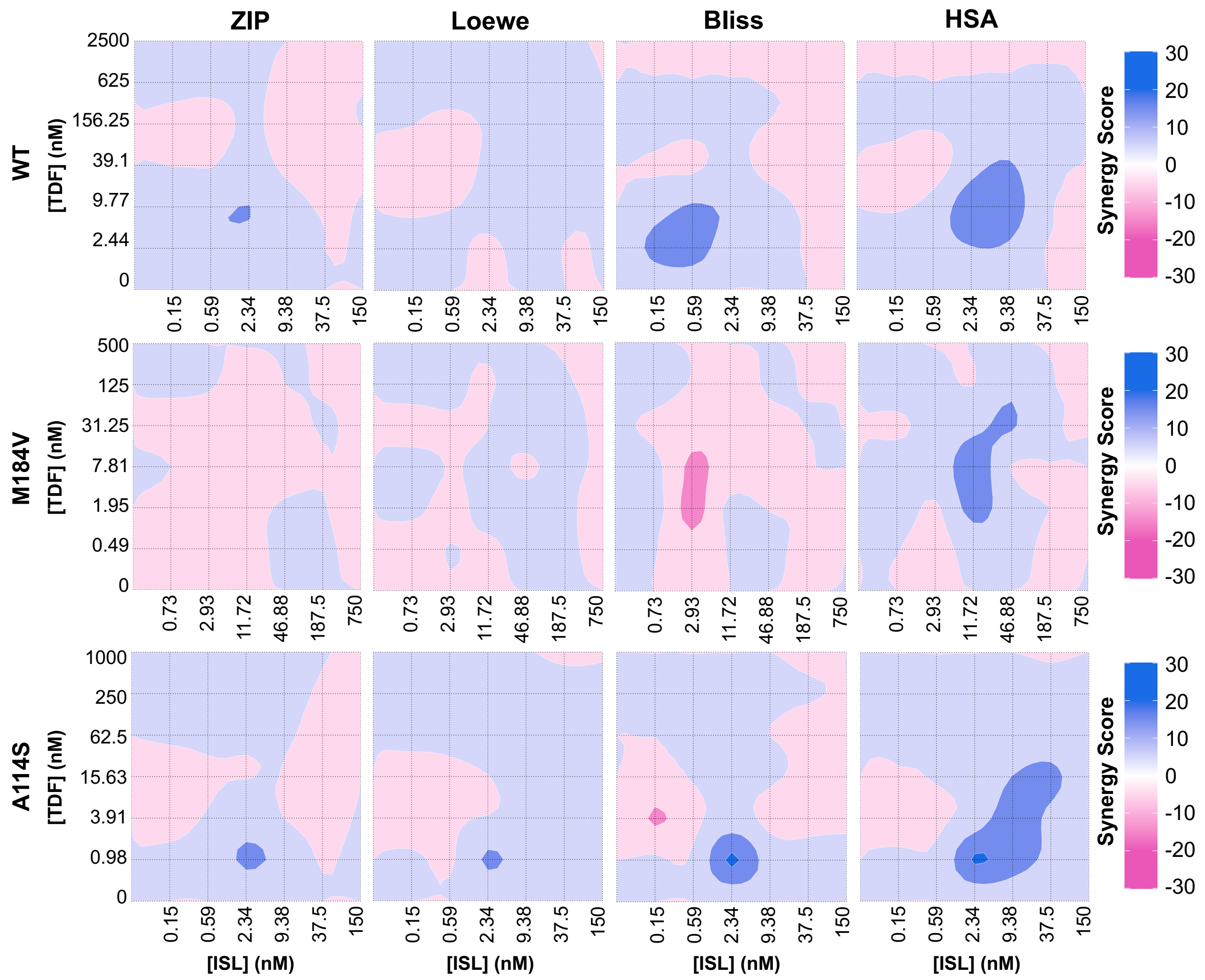


**Fig. S13. Synergy model-based maps of ISL and TDF across VSV-G pseudotyped viral variants.** Maps calculated from three experiments correspond to the ZIP, Loewe, Bliss, and HSA synergy models (left to right). Synergy maps were plotted for -30 ≥ Synergy Score ≥ +30. Color palette: synergism: blue; additivity: white; antagonism: pink. Synergy maps created in SynergyFinder+.
